# The evolution of genetic drift over 50,000 generations

**DOI:** 10.64898/2026.01.25.701616

**Authors:** Joao A. Ascensao, QinQin Yu, Oskar Hallatschek

## Abstract

Random variation in reproductive success–genetic drift–profoundly shapes genetic diversity and evolutionary trajectories. The strength of drift depends on the variance in descendant number, 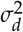, which governs key evolutionary outcomes: for instance, the establishment probability of a beneficial mutation scales inversely with 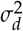. However, whether 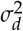 itself evolves over long timescales has remained unclear, because allele-frequency fluctuations depend on drift only through the effective population size, 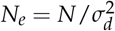, which blends census population size with descendant-number variance. Here, we disentangle these components by using model-based Bayesian inference combined with joint tracking of (i) frequency fluctuations of neutrally barcoded lineages and (ii) census population sizes across growth cycles in the*E. coli*Long-Term Evolution Experiment (LTEE). Analyzing 33 clones spanning the ancestor through 50,000 generations in two replicate populations (Ara-2 and Ara+2), we find that the strength of genetic drift evolved markedly–and divergently–between the two replicate populations. Both census size and 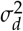 changed substantially through time, with most variation in *Ne* driven by shifts in 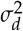 rather than census size. After approximately 2,000 generations, the 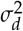 of the two populations diverged sharply: Ara+2 generally remained close to a bottleneck-only null expectation, whereas Ara-2 exhibited 1.5–5× stronger drift, consistent with an evolved increase in stochasticity during growth. These results imply that mutations can alter both the mean and the variance of the descendant-number distribution, and we show that the joint distribution of mutational effects on fitness and on 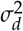 can substantially modulate the rate of adaptation. The key parameter governing genetic drift can therefore itself evolve in the LTEE, with direct consequences for adaptation.

## Introduction

Genetic drift is a stochastic evolutionary force that arises from random variation in reproductive success^1–3^. When drift is strong, it can erode neutral diversity^4^ and accelerate the accumulation of deleterious mutations through Muller’s ratchet^5,6^. Drift also matters in settings where adaptation is common, because it controls the rate at which a newly arising beneficial mutation escape early stochastic loss and establish in the population^2,7,8^ .

For many evolutionary questions, drift is governed primarily by the variance of the descendant-number distribution, 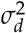. This is evident from standard branching-process approximations for the fate of a weakly beneficial mutation: if a single mutant has mean descendant number1 +*s* with *s*≪ 1, its establishment probability satisfies^2,7,9^ 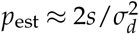. Consequently, doubling 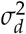 halves the chance that a mutation of given effect *s* establishes, implying that such mutations must arise about twice as frequently to maintain the same establishment rate. If mutations can change 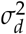 as well as *s*, then, all else being equal, evolution will enrich for mutations that lower the descendant number variance. By contrast, for processes in which drift accumulates over many generations, higher moments of the descendant-number distribution typically contribute little. Quantifying drift therefore largely amounts to measuring the descendant-number variance.

Descendant-number stochasticity has several observable consequences, most directly fluctuations in allele frequencies. For a neutral allele at frequency *f*, the variance of the frequency change per generation is approximately *f*(1 −*f*)/*N* _*e*_, where the effective population size^3^ is 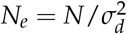. Frequency fluctuations therefore probe drift only through *N*_*e*_, a compound parameter that entangles census size *N* with descendant-number variance 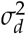. The same fluctuation magnitude can thus arise from a small population with modest reproductive variance or from a large population with highly variable reproduction. Inferring 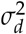 from frequency trajectories therefore requires independent information about*N*.

The concept of effective population size has a complex history in population genetics, with different conceptualizations arising from different modeling assumptions and frameworks^3,10–12^. In a strictly neutral setting over long timescales, *N*_*e*_ determines typical coalescence times (in units of generations) and hence scales with the pairwise diversity^13,14^. However, in populations experiencing strong selection, patterns of genetic diversity reflect a combination of drift and “genetic draft”–the stochastic hitchhiking of alleles linked to selected variants^15–17^. Under such conditions, coalescence times and pairwise diversity can have vastly different relationships to the census population size and forward-in-time descendant number variance^17–19^. Indeed, it has long been noted that effective population size can have very little relation to the census population size^20–23^. This has led to calls for caution in using diversity-based *N*_*e*_ estimates to infer population sizes or predict evolutionary dynamics, especially in adapting populations^11,16^.

Here, rather than inferring genetic drift indirectly from patterns of sequence diversity, we directly measure the short-term descendant number variance by tracking frequency fluctuations of neutral genetic markers over short timescales. By observing how the frequencies of thousands of neutrally barcoded lineages fluctuate from one growth cycle to the next, and measuring census population sizes, we can estimate both 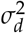 and *N* independently, without relying on assumptions about selection or coalescent structure.

We apply this approach to two replicate populations from the*Escherichia coli*Long-Term Evolution Experiment (LTEE), tracking how the strength of genetic drift has changed over 50,000 generations of evolution. The LTEE was initiated by dividing a single*E. coli*strain into twelve replicate cultures, which were then propagated in a constant environment through daily 1:100 dilutions^24^ (Figure 1A). In this setting, the natural unit of time is the growth cycle, and we define 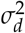 as the variance in descendant number from one dilution to the next for cells sampled immediately after a bottleneck. Random sampling at the bottleneck contributes a baseline variance of one to 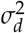; any excess variance reflects additional stochasticity during growth and reduces the effective population size below the bottleneck size, *N*_*b*_. Although the establishment probability of a new mutant depends on when during the cycle it arises because population size changes over time^9^, the*rate*of establishment is governed by the classical expression 2*sN*_*e*_*µ*, where*µ*is the mutation rate and*s*is the per-cycle excess in relative growth (see Supplementary NoteS2).

**Figure 1.**
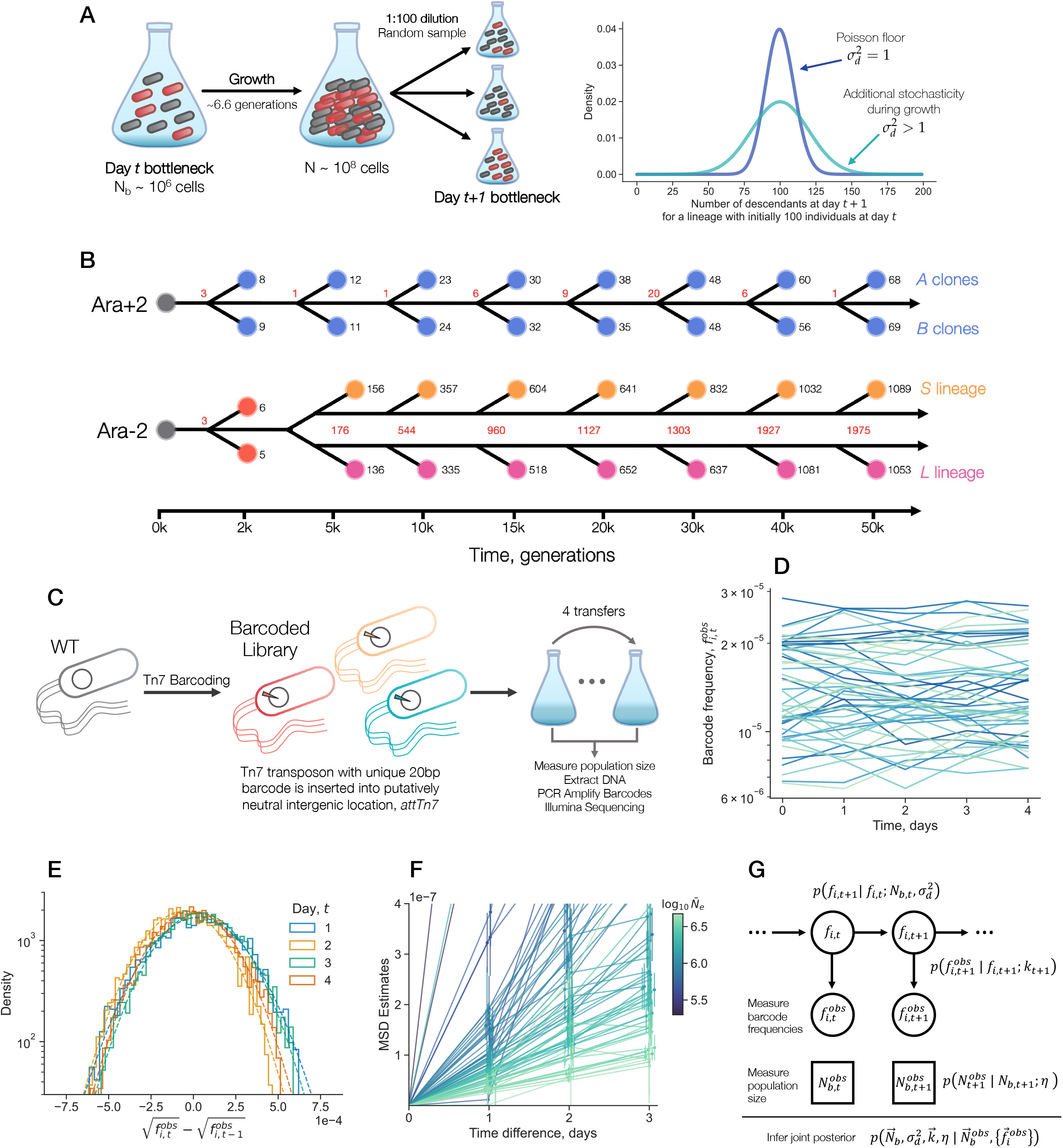
Measuring the strength of genetic drift over evolutionary time. (**A**) Representation of the growth and dilution process in the LTEE. We track two neutral lineages, shown as black and red cells. Cycle*t*begins after a bottleneck. Over 24 hours, the population grows 100-fold, undergoing approximately 6.6 generations. At the end of the cycle, a 1:100 dilution is transferred to fresh media. Because of this sampling step, the number of descendants a lineage contributes from one cycle to the next is a random variable with at least Poisson variance; any additional variance implies another source of stochasticity in the growth process. (**B**) Schematic of the phylogenies and clone selection for the two LTEE lines examined in this study, Ara+2 and Ara-2. We sampled two clones from each timepoint in each line. The Ara-2 line diversified into the*S*and*L*lineages early in the experiment, while the Ara+2 line did not diversify into multiple long-lasting coexisting lineages. Black numbers represent the total number of mutations relative to the ancestor for each clone; red numbers represent number of different mutations between clones at the same time point. (**C**) We generated many neutral variants of each LTEE clone by using a Tn7 transposon system to insert random barcodes at a single intergenic locus. The resulting libraries were passaged under LTEE conditions and sampled daily. (**D**) Example barcode frequency trajectories, from the LTEE ancestor (REL606) library (replicate 1). (**E**) Transition densities of barcode frequencies for the REL606 library, replicate 1. Dotted lines represent Gaussian fits to the empirical densities. (**F**) Estimates of the mean squared displacements (MSD) of the square-root transformed barcode frequencies. Line color indicates the estimated 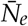; error bars represent 95% confidence intervals. (**G**) Statistical model and inference strategy for estimating genetic drift parameters. The unobserved “true” barcode frequencies *f*_*i,t*_ evolve according to a specified transition density, and we additionally model the technical noise from measuring barcode frequencies and total population size. Given the data, we estimate the joint posterior of all parameters via MCMC.

We focus on two LTEE populations–Ara-2 and Ara+2–that represent distinct eco-evolutionary outcomes: Ara-2 diversified into two coexisting ecotypes around 6,500 generations and evolved hypermutability^25–28^, while Ara+2 retained the ancestral mutation rate and did not form a stable ecological structure.

We isolated 33 clones from the two replicate populations of the *E. Coli* Long-Term Evolution Experiment, spanning the ancestral strain through 50,000 generations of evolution. For each clone, we generated thousands to tens of thousands of neutrally barcoded variants and tracked their frequency fluctuations over several days of growth. Using a Bayesian inference framework applied to barcoded lineage trajectories, we find that both the census population size and descendant number variance change substantially through evolutionary time. Notably, while the census population sizes of the two lineages follow similar trajectories, their descendant number variances diverge markedly after approximately 2,000 generations. Taken together, our findings indicate that mutations can modify not only the mean but also the variance of the descendant-number distribution. We demonstrate that the joint distribution of mutational effects on fitness and on 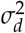 can appreciably tune the rate of adaptation. Our results demonstrate that the descendant number variance is a highly evolvable trait in LTEE populations, with direct consequences for adaptation.

## Results

### Measuring genetic drift parameters

To map out the evolution of genetic drift in the Ara-2 and Ara+2 lines, we first isolated sixteen clones from each line, from eight timepoints spread between two thousand and fifty thousand generations of evolution (Figure 1B). For Ara+2, we used the previously isolated^29^ *A*and*B*clones from each time point; the labels were arbitrarily assigned and do not reflect phylogenetic structure. For Ara-2, we used the *A* and *B* clones from 2k generations. Around 6.5k generations, the Ara-2 population split into two coexisting lineages that have coevolved for the duration of the experiment–*S*and*L*. Thus, for all timepoints 6.5k generations and after, we used one *S*clone and one*L* clone. We also included an LTEE ancestor (REL606) in our study. We sequenced the genomes of the four clones that had not previously been sequenced. We see that both lines accumulate mutations over time relative to the REL606 ancestor; the Ara-2 lineage has a much faster mutation accumulation rate due to its hypermutator phenotype (Figure 1). As expected, we see that the number of mutations separating the *S*and*L* clones increases over evolutionary time, while the genetic distance between Ara+2*A* and *B* clones does not systematically change.

We sought to measure the strength of genetic drift–captured by the values of parameters 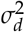, *N*_*b*_, and *N*_*e*_–for each of the 33 LTEE clones. Our approach to measure the three parameters is based on observing the frequency trajectories of many neutral variants of a clone, as the frequency fluctuations are related to the strength of genetic drift. For neutral lineages that are composed of sufficiently many individuals such that the central limit theorem applies, the lineage frequency at time *t*+1,*f* _*t*+1_, can be written as the following Markov chain,

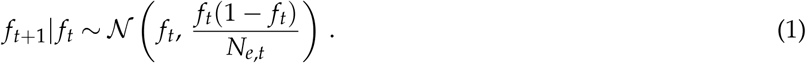

We measure lineage frequencies at the end of the 24-hour growth cycle, effectively coarse-graining over the within-cycle dynamics. Accordingly, each time step corresponds to one dilution-to-dilution interval, and *σ*^2^ denotes the bottleneck-to-bottleneck variance in descendant number for cells sampled immediately after dilution. We only consider small lineage frequencies, *f*_*t*_ ≪1, so that we can ignore the factor of (1− *f* _*t*_) in the variance. We allow the effective population size to vary in time, as we have previously observed large abundance fluctuations in LTEE clones^30^, so it is necessary to account for potentially changing population sizes. We assume that the descendant number variance stays approximately constant over the timecourse, so we can write the (potentially time-dependent) effective population size as,

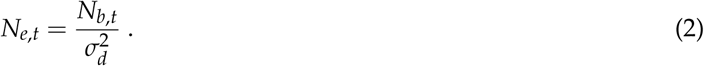

Thus, we must measure the bottleneck population size in addition to the frequency trajectories to isolate the contribution of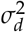 to *N* _*e,t*_.

To experimentally generate many neutral variants of each LTEE clone, we used a previously developed method^31^ where random 20bp barcodes are inserted into a putatively neutral genomic location,*attTn7*, by a Tn7 transposon system (Figure 1C). We aimed for approximately 1,000-10,000 barcodes per strain, which we found was a sufficient number to accurately resolve the magnitude of frequency fluctuations, while allowing us to sequence many experiments on a single Illumina lane. To measure the frequency of each neutrally barcoded variant, we amplified the barcode region via PCR (polymerase chain reaction), and used Illumina amplicon sequencing to count the number of times each barcode appears (Figure 1D). We propagated the Tn7 barcoded libraries for each clone in the LTEE evolutionary condition (without any other strains present in culture), sampling the population every day to (i) extract genomic DNA for later barcode sequencing and (ii) measure the bottleneck population size by counting colony forming units (CFUs). For each strain, we performed two biological replicate experiments, on different sets of days.

Following barcode sequencing, we quantified barcode reads and performed error correction on the sequences. Subsequently, we filtered out barcodes in each time course that appeared to have non-neutral fitness effects (relative to the other barcodes), putatively caused by mutations that arose on the background of the barcoded lineage. Generally, only a small percentage of barcodes displayed non-neutral trajectories. We simulated the data generating process and computational pipeline to ensure that the filtering process did not inadvertently cause biases or other pathologies to appear (Figures S9-S13).

As we are measuring barcode frequencies by counting sequencing reads, there must be measurement error in the frequencies that is at least as strong as Poisson noise; any additional noise in the sequencing process will cause overdispersion. But again, if the read counts are sufficiently large, then the central limit theorem applies, and we can model the measurement error as Gaussian,

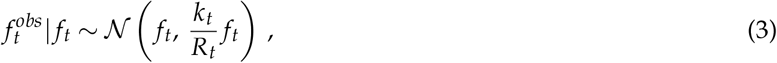

where *k*_*t*_ is the measurement noise overdispersion, and *R*_*t*_ is the total number of sequencing reads. We allow *k*_*t*_ to vary over time because it has been previously shown that overdispersion varies across samples in similar barcoded high-throughput sequencing-based experiments^32,33^, presumably caused by fluctuations in e.g. PCR conditions or genomic DNA extraction efficiency. The two overdispersion parameters, 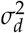 and *k*_*t*_, are structurally identifiable in our model. Intuitively, this is because the variance due to descendant number variance increases over time, while the measurement noise is uncorrelated in time.

We use a simple square-root transformation on both the true and observed frequencies as a variance stabilizing transformation^34^; i.e. the square-root transformed frequencies still have Gaussian densities, but the variance is no longer dependent on the mean. The observed transition densities result from a convolution between genetic drift and measurement noise. Using technical replicates, we show that the measurement noise of square-root transformed frequencies is very well fit by a Gaussian (Figure S6). Together, this would imply that observed transition densities between the square-root transformed observed frequencies at neighboring times should follow a Gaussian distribution, providing a check on our model. Indeed, we see that the observed transition densities are very well fit by Gaussian densities, well into the tails of the distribution (Figure 1E).

Our model also predicts that variance in a barcode frequency trajectory should increase over time (Figure S1). Specifically, we expect that the mean squared displacement (MSD) between square-root transformed barcode frequencies should increase linearly over time, with a slope proportional to 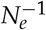,

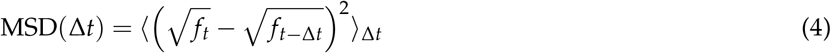

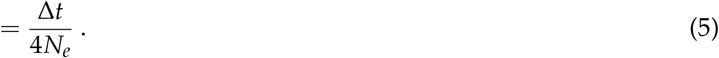

This simple relationship assumes that *N*_*e*_ is constant over time, and does not include the effects of measurement noise. However, we were able to use a simple procedure to account for measurement noise (see Methods), and obtained MSD estimates for the barcoded libraries (Figure 1F). We see that the MSD across strains and experiments generally increases approximately linearly over time (Figure 1F); slight deviations from linearity are likely due to the well-known sensitivity of empirical MSDs to measurement noise^35–37^.

In order to more quantitatively estimate the genetic drift parameters for each time series, we developed a Bayesian inference approach. Following Yu et al. (2024)^34^, we combine the likelihoods of equations 1 and 3, and integrate over the hidden “true” barcode frequencies to define a Kalman Filter (continuous-state hidden Markov model) of the observed barcode frequencies (Figure 1G). We also define a likelihood to quantify the measurement error of the observed bottleneck population size, 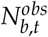. We combine the likelihoods with non-informative/weakly informative priors, and use a Hamiltonian Monte Carlo (HMC) approach to sample from a joint posterior of all the parameters (see Methods).

### The strength of genetic drift varies across evolutionary time

We obtained joint posterior estimates for all of the genetic drift and measurement noise parameters, across all experiments and biological replicates. We show representative examples of marginal posteriors in Figure 2. Here and throughout, 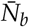 and 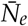 represent the harmonic mean over experimental timepoints of the bottleneck and effective population size, respectively; the harmonic mean is the evolutionarily relevant parameter^38^. We see that the bottleneck population size is generally not constant over time, or across biological replicates (Figure 2C), consistent with prior experiments that have observed large abundance fluctuations in LTEE populations^30^. As an example, we see that the 10k *S* and *L* clones show consistently different effective population sizes and descendant number variances (Figure S2). Additionally, the sequencing measurement error overdispersion, *k*_*t*_ is oftentimes close to one–indicating noise dominated by Poissonian sampling–but the magnitude is sometime uncertain, and sometimes almost certainly greater than one (Figure 2C).

**Figure 2.**
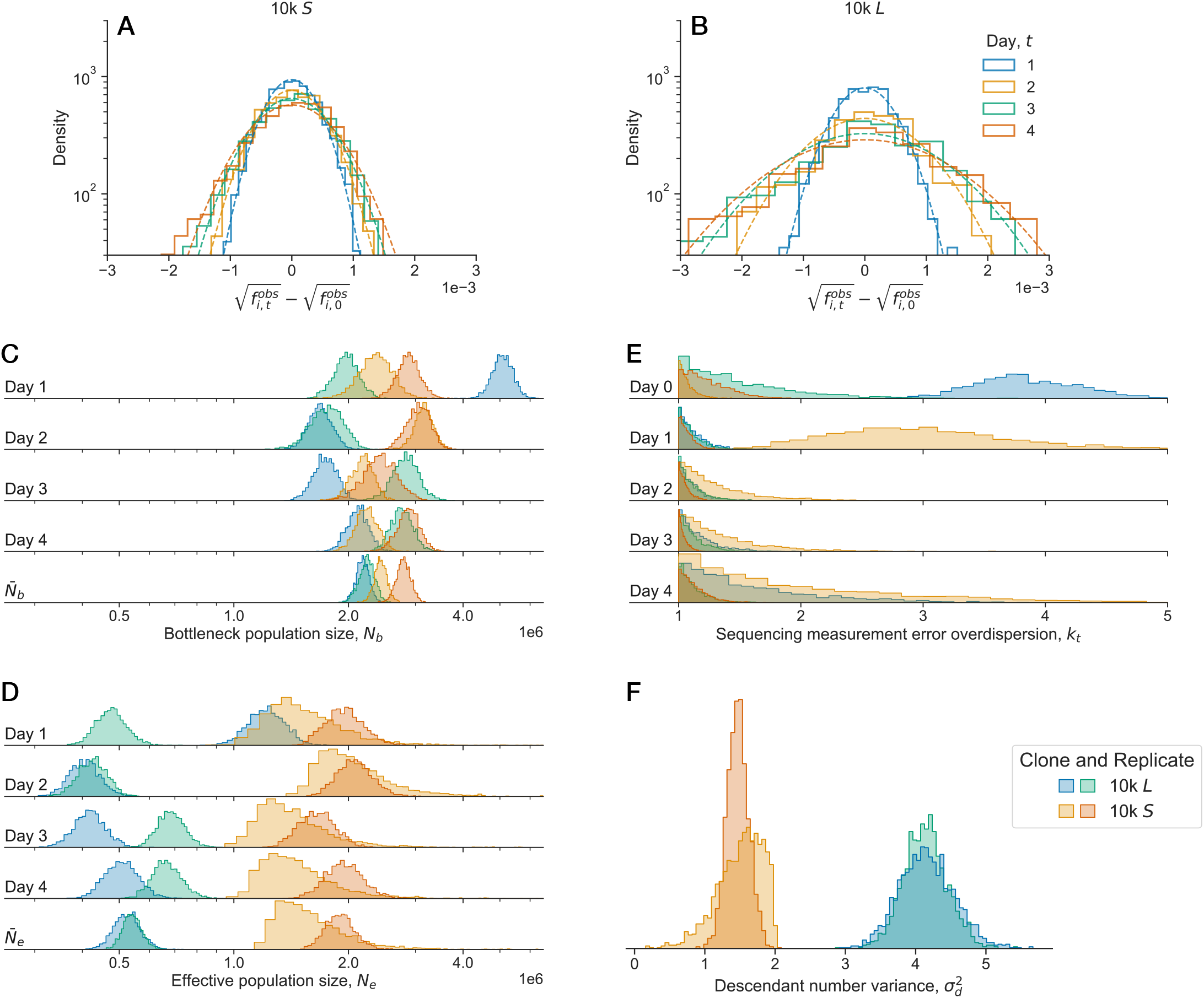
Examples of transition and posterior densities. Data from the*S*and*L*clones sampled from 10k generations. (**A-B**) Empirical transition densities of barcode frequencies; dotted lines represent Gaussian fits to the empirical densities. Here, we compute the transition densities in reference to the frequency at time point 0. As suggested by the estimates of *N*_*e*_, the neutral barcode trajectories of the *S* clone accumulate less variance than those of the *L* clone. (**C-D**) Posterior estimates of genetic drift parameters; different colors represent different libraries and biological replicates. Posterior estimates of the (**C**) bottleneck and (**D**) effective population sizes over the experimental time points. We also show the harmonic mean of the population sizes over the four experimental time points, 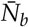 and 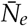. Posterior estimates for the (**E**) sequencing measurement error overdispersion and (**F**) descendant number variance.

We then systematically compared the genetic drift parameters–the bottleneck and effective population size, and the descendant number variance–across biological replicates and strains (Figure 3). As the bottleneck and effective population sizes are allowed to vary over time, we focus on the harmonic mean across time points for both parameters. The correlation between biological replicates is generally quite good and greater than zero for all three parameters, ranging from about 0.65 to 0.9 (Figure 3A-C).

**Figure 3.**
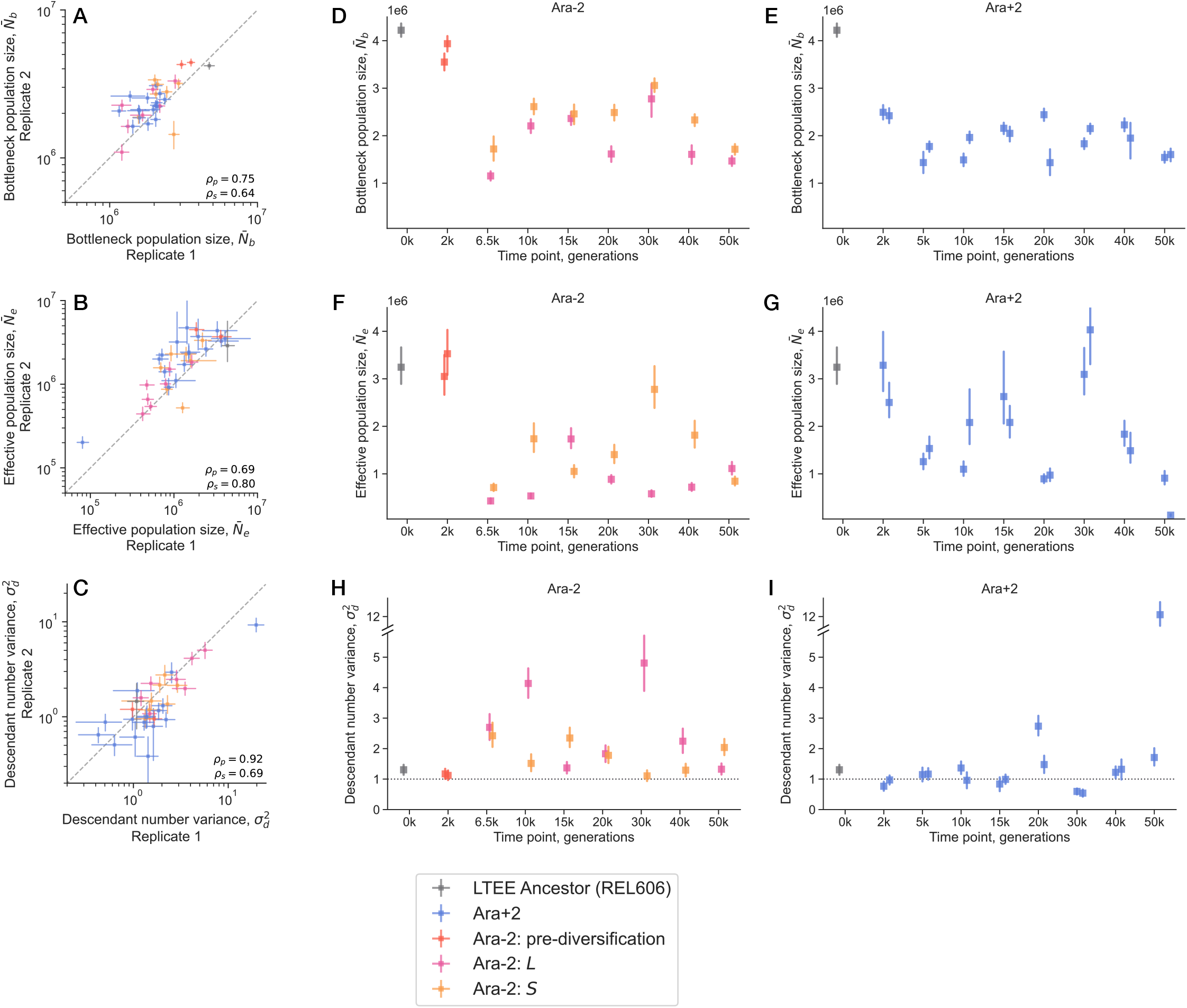
Estimates of genetic drift parameters over evolutionary time. (A-C) Correlations of inferred parameters between biological replicates; *ρ*_*p*_ and *ρ*_*s*_ represent Pearson’s and Spearman’s correlation coefficient, respectively. (D-I) Estimates of the inferred parameters, for each sampled clone, over evolutionary time. All point estimates represent posterior medians, and error bars represent 95% credible intervals.

We combined the data of the two biological replicates for each strain to obtain final posterior estimates for all parameters. We see that the LTEE ancestor has a mean bottleneck population size around4 · 10^6^, which is the highest among all strains (Figure 3D-E). This is consistent with the fact that nearly all MSD lines are higher than the ancestor (Figure 1F). We observe that the mean bottleneck population size quickly starts to decrease as evolution proceeds, consistent with prior measurements^39^. The rate of decrease is initially faster in Ara+2 compared to Ara-2–at 2,000 generations, the Ara-2 clones had mean bottleneck population sizes around3.5 · 10^6^, while the Ara+2 clones were around2.5 ·10^6^. After 2k generations, the bottleneck population size fluctuates around1 −2 · 10^6^ for clones in both lines. The effective population size also varies widely over the course of evolution, but notably it has markedly distinct dynamics compared to the bottleneck population size (Figure 3F-G). This difference is caused by the complex changes in descendant number variance over evolutionary time.

The changes in the descendant number variance over time shows stark differences between the Ara+2 and Ara-2 lines (Figure 3H-I). For both lines, we see that the descendant number variance may be declining slightly between zero and two thousand generations. The LTEE ancestor has a descendant number variance around 1.3, indicating that while the primary source of variance is indeed the once-per-cycle bottlenecks, there is some additional unexplained variance, putatively caused by stochasticity in the growth process. All four clones at 2k generations have a descendant number variance indistinguishable from 1. After two thousand generations, the Ara+2 and Ara-2 lines diverge sharply. The majority of clones in Ara+2 show descendant number variances that hover around one, the exceptions being the clones at the 20k and 50k timepoints. All four clones in those timepoints display descendant number variances significantly above one. The 50k*B*clone shows especially outlying behavior, with a *σ*^2^ about 12 fold larger than expected from bottlenecking alone; the excess strength of fluctuations can be visually seen in the barcode trajectories (Figure S2). In Ara-2, the descendant number variance sharply increases immediately following diversification into the*S*and*L*lineages. At 6.5k generations (immediately after diversification), both*S*and*L*show descendant number variances around 2.5, around twice as large as the ancestor.

It may be that the initially similar descendant number variances of*S*and*L*is due to the fact that they have not yet had the chance to significantly genetically diverge. After 6.5k generations, the descendant number variances start to become more uncoupled from each other, varying significantly over evolutionary time, while almost always staying larger than one.

### Quantifying systematic differences in the evolution of genetic drift between lineages

From the marginal genetic drift posteriors, we saw systematic differences in the evolution of genetic drift between Ara+2 and Ara-2. We sought to quantify the magnitude of such differences, so we obtained posteriors for several summary statistics across clones in each LTEE lineage (Figure 4).

**Figure 4.**
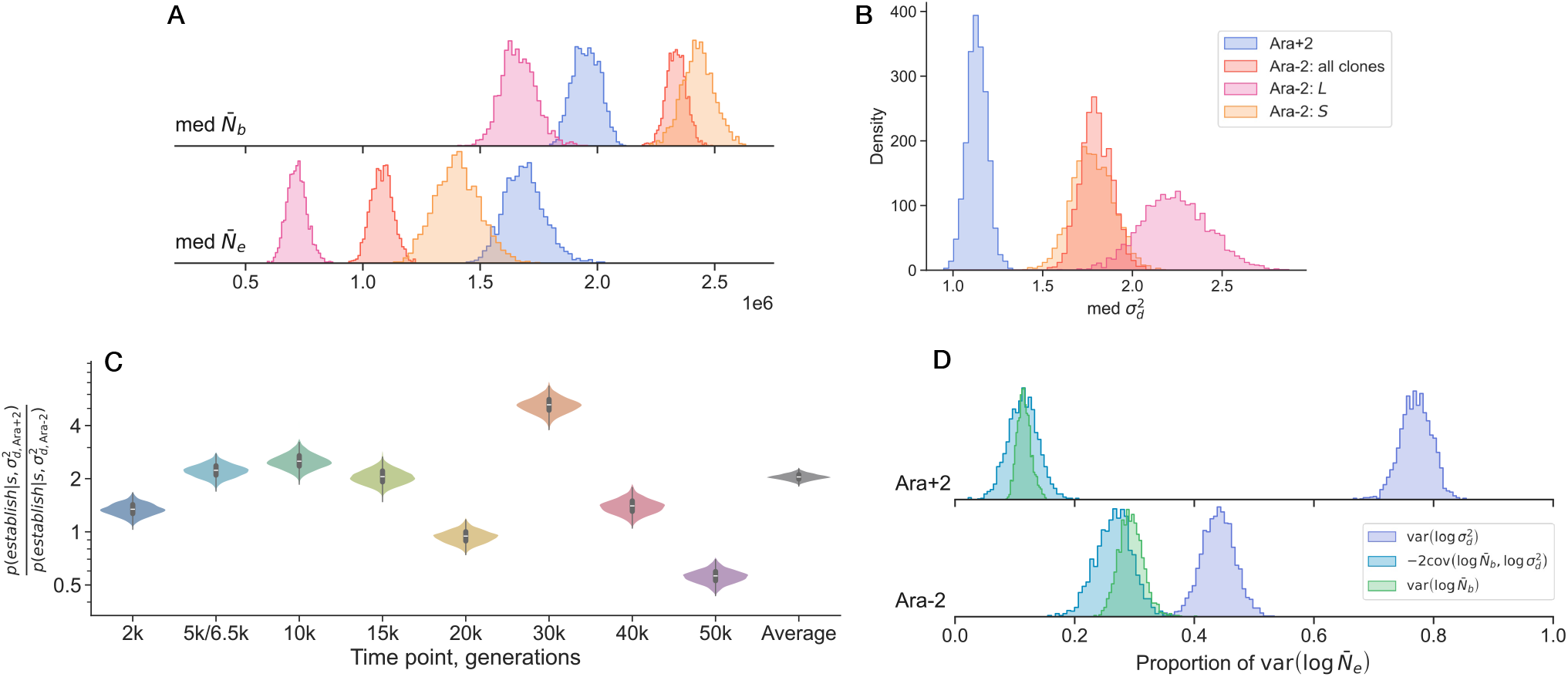
Patterns and consequences of the strength of genetic drift across lineages. (**A-B**) Posterior density estimates of the median (**A**) bottleneck and effective population sizes and (**B**) the descendant number variance, over all clones within a group. (**C**) Posterior densities of the ratio of establishment probabilities, *p*_*est*_(Ara+2)/*p* _*est*_(Ara-2), for a beneficial mutation of given fitness effect*s*arising on an Ara+2 versus an Ara-2 background; values greater than one indicate that the mutation is more likely to establish in Ara+2. For each time point, we computed this ratio for all four pairings of one Ara+2 clone with one Ara-2 clone, and averaged across the four pairs. The 6.5k clones from Ara-2 were compared to the 5k clones of Ara+2. The rightmost density (“Average”) shows the mean ratio across all timepoints. (**D**) Posterior densities of the contributions to the variance of log effective population sizes, across clones in a lineage.

As a simple first question, we wished to compare the typical bottleneck and effective population size across Ara-2 and Ara+2. To that end, we computed the posterior of the median 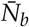 and 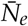 across all clones within a line (Figure 4A), i.e. *p*(med_*i*_ 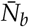) and *p*(med_*i*_ 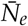). We see that typically, 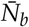 is higher in Ara-2 clones compared to Ara+2 clones, by a factor of about 25%. However, once we separate the Ara-2 by ecotype lineage, we see that*L*clones actually typically have the lowest 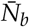, while*S*clones have the highest. The pattern changes when we shift our attention to the effective population size. In contrast to the case with 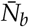, the Ara+2 clones typically have larger 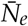 than all Ara-2 clones, as well as when compared to the*S*and*L*clones (when the lineages are separated). This discrepancy between the typical bottleneck and effective population sizes between the LTEE lines is due to the fact that Ara-2 clones typically have a larger descendant number variance compared to Ara+2 clones (Figure 4B). We see that the typical 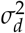 for Ara+2 clones is around 1, while it is above 1 for Ara-2 clones. In a slightly different way of quantifying differences in descendant number variance, we computed the proportion of clones with 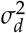 larger than some cutoff value (Figure S3A). We see that nearly all Ara+2 clones have a descendant number variance lower than 1.5, while the Ara-2 clones span a larger range of values. Together, these observations underscore how differences in effective population size between populations can be decoupled from their relative census population sizes by factors of order 1.

The differences in descendant number variance between Ara+2 and Ara-2 have direct implications for the fate of beneficial mutations. Because the establishment probability of a weakly beneficial mutation scales as 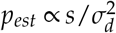, a mutation of given fitness effect s will establish at different rates depending on the genetic background on which it arises. To quantify this, for each timepoint we computed the ratio of establishment probabilities, *p*_*est*_(Ara+2)/*p* _*est*_(Ara-2), for every pair of clones across the two populations, and averaged over the four pairs (Figure 4C). Averaging across all timepoints, a beneficial mutation of given effect is roughly 2-fold more likely to establish in Ara+2 than in Ara-2. This advantage, however, is far from constant over evolutionary time. The ratio is modest at 2k generations (∼1.3), when 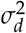is similar in the two lines, and grows substantially after Ara-2 diversifies, reaching ∼2.2 at 6.5k, ∼2.5 at 10k, and ∼2.1 at 15k generations. The disparity peaks at 30k generations, where mutations are roughly 5-fold more likely to establish in Ara+2 than in Ara-2. At 20k and 40k generations the ratio is closer to one, and notably at 50k the ratio inverts (∼0.56), reflecting the outlying *σ*^2^ of the Ara+2 50k B clone (Figure 3I); at this timepoint, beneficial mutations are predicted to establish more readily in Ara-2. Overall, the establishment probability of a beneficial mutation of given effect can differ by factors ranging from <1 to ∼5between the two populations, depending on the evolutionary timepoint–a substantial and time-varying bias that arises purely from differences in the strength of genetic drift, independent of any differences in the underlying distribution of fitness effects.

We observed that the effective population size varied strongly over evolutionary time, for both the Ara+2 and Ara-2 lines (Figure 3F-G). However, as 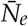 is a compound parameter of 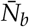 and 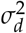, it is not immediately clear which constituent parameter causes most of the variance in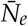 across clones. To assess the contribution of the two parameters to 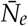, the variance of log 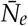 across all clones in a line can be decomposed into three components,

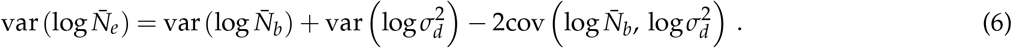

We thus compute the three components across all clones within a line, and normalize each component to var(log 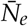) (Figure 4E). We focused on log 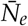 as it has a simple additive relationship to the bottleneck population size and descendant number variance. We see that variance in log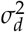 has the largest contribution to variability in log 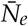 across clones, for both lines. A portion of this pattern is a result of the outlying 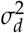 of the Ara+2 50k*B*clone, and the relatively large 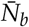 of REL606; however, even when we excluded either or both of those clones from our analysis, we saw the same qualitative pattern (Figure S4). This analysis suggests that most of the variability in *N*_*e*_ across evolutionary time arises from changes to the descendant number variance, not from changes to the census population size.

### Causes and consequences of 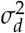evolution

We hypothesize that noise in growth traits could give rise to additional effective descendant number variance. Previous studies have documented stochastic variability in both the exponential growth rate and lag time among sister microbes grown in the same environment^40–44^, providing a concrete mechanism by which additional stochasticity during growth can manifest in well-mixed, genetically homogeneous populations. To investigate this, we used simulations and analytic theory to study asexual growth processes with noise in the doubling time and lag time, and connected such noise to effective additional descendant number variance (Figure 5A-B, SI sectionS3). Both stochastic lag and doubling times can in principle contribute to increases in 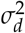. A key feature of our theoretical results is that 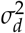depends on both the *mean* and the *variance* of the underlying lag and doubling-time distributions: even a mutation that shifts only the mean lag or doubling time, leaving cell-to-cell variability unchanged, will generically alter 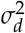. We applied our theoretical results to predict how previously measured distributions of lag^40^ and doubling times^41^ in non-LTEE*E. coli*strains would impact the descendant number variance in a serial dilution environment. If the measured*E. coli*strains are not too different from LTEE strains, then we would conclude that noise in lag time is predicted to produce a much larger increase in 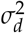 than noise in doubling time.

**Figure 5.**
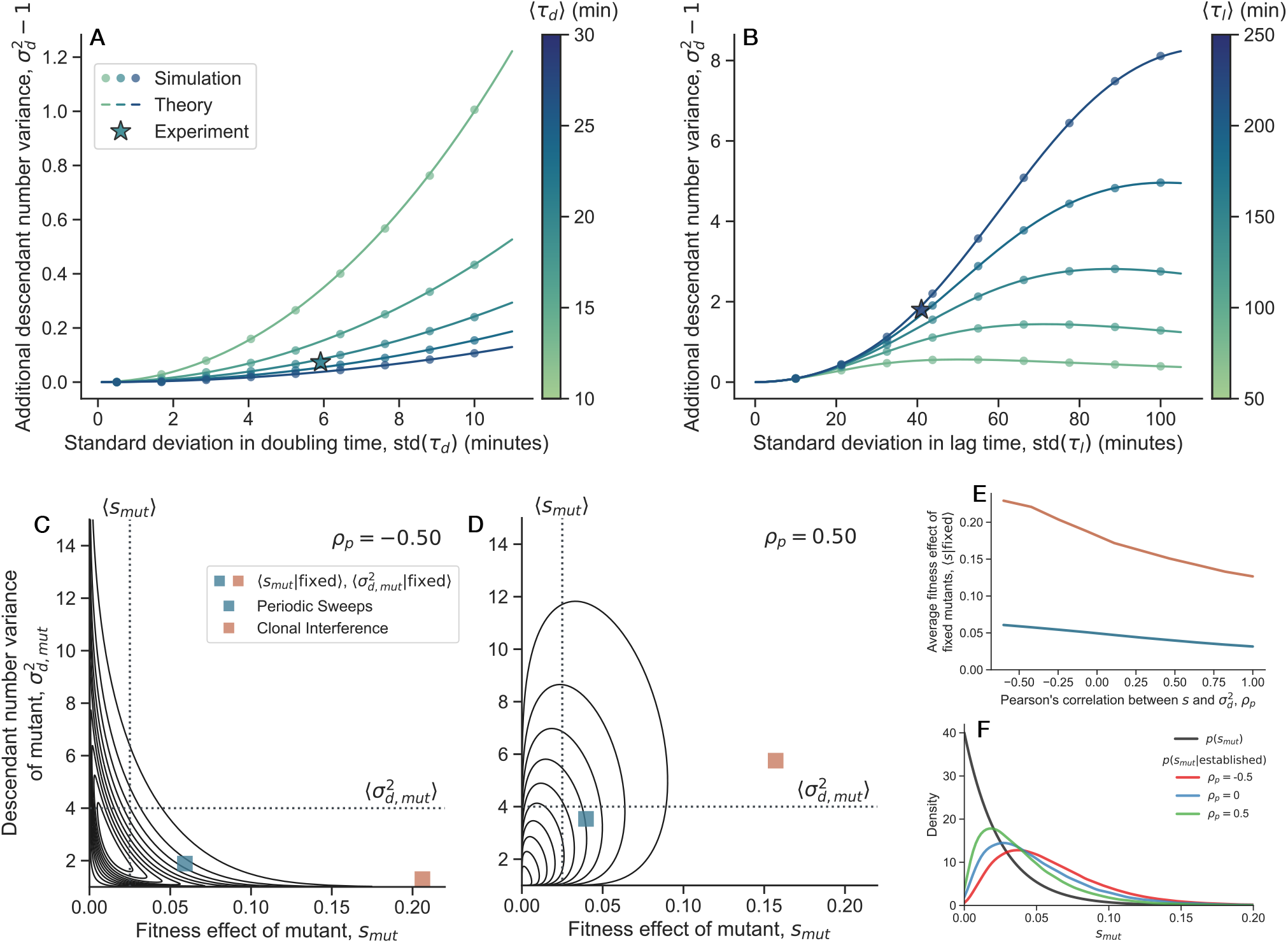
Possible physiological and evolutionary causes and consequences of descendant number variance evolution. (**A-B**) Additional contributions to the descendant number variance beyond that from bottlenecks can arise from noise in (**A**) the exponential doubling time,*τ* _*d*_, or (**B**) the time in lag phase,*τ* _*l*_ . We model both*τ* _*d*_ and*τ* _*l*_ as gamma random variables, simulating the stochastic growth process (circle symbols) and obtaining analytical predictions (SI sectionS3). We use the theoretical results to predict the additional descendant number variance that would arise from previously experimentally measured distributions of doubling times^41^ and lag times^40^ in non-LTEE strains of *E. coli*(star symbols). Note that the axes for each panel are on different scales. (**C-D**) We hypothesize that for any given genetic background, there is a joint distribution of mutant fitness effects and mutant descendant number variances,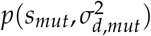)(black contour lines). Here, both marginal distributions are exponential and linked with a Gaussian copula; contour levels correspond to iso-proportions of the density. Dotted lines represent marginal means of the distribution; marginal distributions are held constant as correlation is varied. When an adaptive mutant fixes in the population, it can change both the fitness and descendant number variance. The probability of mutant fixation depends on both the mutant’s fitness effect and descendant number variance. Evolutionary dynamics simulations demonstrate that on average, adaptation will bias fixed mutants towards larger fitness effects, and smaller descendant number variance (square symbols). However, the magnitude and direction of such biases is affected by both the evolutionary regime (periodic sweeps versus clonal interference) and the correlation structure of 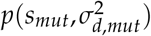. (**E**) In both regimes, a stronger positive correlation between *s*_*mut*_ and 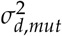 is associated with a lower ⟨*s*_*mut*_|fixed⟩. (**F**) We compare the distribution of fitness effects of new mutations (black) to the distribution of fitness effects of*established*mutations (color). Correlations between *s*_*mut*_ and 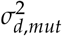
systematically deforms the distribution of established mutations. Here,*ρ* _*p*_ represents Pearson’s correlation coefficient of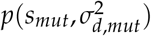.

Does 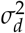 evolution affect the rate of adaptation, and what evolutionary forces might cause 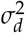 to change over time? There are a number of possible ways in which 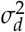 can affect evolutionary dynamics. (1) Theory predicts that differences in 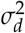 between two otherwise neutral variants would generate quasi-selective effects favoring the variant with lower 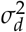, although this effect is likely weak because the effective selective effect scales as1 /*N* . (2) The descendant number variance can directly affect the rate of adaptation. In the periodic sweeps (strong selection, weak mutation) regime, this effect can be substantial, as the rate of adaptation scales inversely with 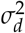. However, in the clonal interference (weak selection, strong mutation) regime, this effect is likely weak, as the rate of adaptation only depends logarithmically on 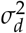, because adaptation is not limited by the supply of established beneficial mutations.(3) In principle, a variant that reduces offspring number variance 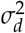 and thus increases the establishment rate of beneficial mutations acts analogously as a mutation-rate modifier. Such variants would be favored by indirect selection on linked beneficial mutants whose establishment they enabled, with the same qualitative population-size and mutation-rate scaling as other mutation-rate modifiers^46,47^. (4) We find that correlations between the the fitness and drift effects of mutations can substantially shape both which mutations fix and the overall speed of adaptation. As noted above, changes in 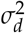may arise as a correlated response to selection on other traits: if selection in the LTEE has acted on mean lag duration or doubling time for reasons unrelated to drift per se (e.g., to optimize use of fresh media), the corresponding shifts in growth-trait distributions will alter 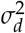 as a side effect. The sign of this correlated response depends on which trait selection targets: because shorter doubling times tend to increase 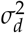 (Figure 5A), selection for faster exponential growth would generate a*positive*correlation between *s*_*mut*_ and 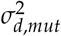 among new mutations, whereas shorter lag times reduce 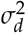(Figure 5B), so selection for shorter lag times would generate a *negative* correlation.

These considerations motivate extending the distribution of fitness effects (DFE) to a joint distribution over mutant fitness effects and descendant number variances, 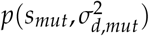 (Figure 5C-E). When a mutant arises, its establishment probability depends on both *s*_*mut*_ and 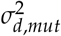 (i.e. the probability that mutations survive drift and start growing exponentially^48^). So, the distribution of *fixed* mutations generally differs from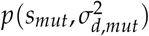. In the periodic sweeps regime, established mutations will almost certainly fix in the population. However, in the clonal interference regime, established mutations still have to compete with each other for fixation. The average effect of an established mutation is,

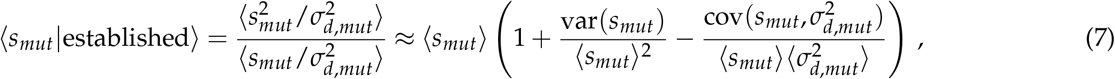

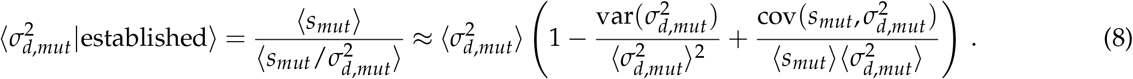

When *s*_*mut*_ and 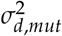are uncorrelated, these reduce to the classical expressions ⟨*s*_*mut*_|fixed⟩=⟨*s* ^2^⟩/⟨*s*⟩ and 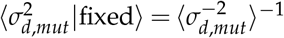deviations from the classical expression for ⟨*s*_*mut*_|fixed⟩ arise specifically from correlation between *s*_*mut*_ and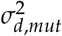.

Crucially, the same qualitative effect persists in the clonal interference regime (Figure 5C-E), where the rate of adaptation depends only logarithmically on 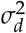 in isolation, but can still be substantially modulated by the correlation between fitness and drift (Figure S14). The intuition is that establishment jointly favors larger *s*_*mut*_ and smaller 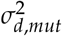, so correlations between the two systematically deform the establishment-weighted DFE relative to 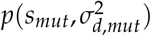: a negative correlation right-shifts the effective DFE, because highly beneficial mutations get an extra boost from being low-drift, while a positive correlation left-shifts it, because highly beneficial mutations are penalized by being high-drift (Figure 5F). In both regimes, stronger positive correlations between *s*_*mut*_ and 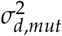 therefore lower ⟨*s*_*mut*_|fixed⟩ and raise 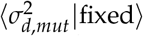 (Figure 5E,S15). Notably, even when *s*_*mut*_ and 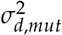 are uncorrelated in the clonal interference regime, the average fitness effect of a fixed mutation still depends on the distribution of 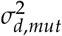 (Figure S17), putatively due to indirect effects such as 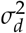-modifying mutations acting as effective mutation-rate modifiers^46,47^. The structure of 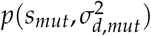–currently unknown for any system–thus likely plays an important role both in shaping the evolution of 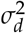and in setting the rate of adaptation, even in regimes where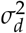 alone has only a weak effect. We present this framework as a hypothesis-generating tool rather than as an explanation for the specific 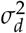 trajectories we observe; testing it would require direct measurements of the joint fitness-drift distribution.

## Discussion

In this work, we measured how genetic drift changed over 50,000 generations in two*E. coli*populations from the Long-Term Evolution Experiment (LTEE). By pairing neutral genetic barcodes with a Bayesian inference framework, we disentangled the two components of the effective population size, *N*_*e*_: the census population size (*N*_*b*_) and the descendant-number variance 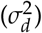. In both populations, *N*_*b*_ and *N*_*e*_ fluctuated substantially through time, with most of the variation in *N*_*e*_ driven by shifts in 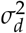. After approximately 2,000 generations, the two populations diverged: their census sizes followed similar paths, but their descendant number variances–and thus the strength of genetic drift–separated markedly. These differences have direct evolutionary consequences in this system: because the establishment probability of a beneficial mutant is 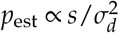, a mutant of a given fitness effect is roughly twice as likely to establish in Ara+2 as in Ara-2 on average across the timecourse.

We emphasize that these conclusions are based on two replicate populations from a single experimental system, propagated under a specific serial-transfer regime with daily 1:100 bottlenecks in a constant minimal-glucose environment. The extent to which 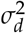 evolves in other systems–natural microbial populations, eukaryotes, or microbial populations under different propagation regimes–remains an open empirical question. The recurring lag phase imposed by daily dilution may make the LTEE particularly conducive to evolutionary changes in 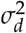, since noise in lag time appears to be a potent contributor to excess descendant number variance (Figure 5B); continuous-culture systems and populations with very different life histories may behave differently. That said, the broader phenomenon of substantial reproductive-number variance unaccounted for by census size is not unique to the LTEE. Highly fecund marine organisms such as oysters and cod exhibit very low *N*_*e*_/*N* ratios attributed to “sweepstakes” reproductive success^49–51^, SARS-CoV-2 transmission shows large temporal variation in *N*_*e*_ driven by fluctuating superspreading^34^, and *Streptomyces* populations exhibit heavy-tailed descendant-number distributions that vary among closely related strains^52^. Our work contributes a direct demonstration that, at least in one system, the magnitude of 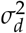 can meaningfully evolve over relatively short evolutionary timescales.

Our findings raise a potential pitfall in interpreting patterns of genomic variation. Many demographic inference methods aim to reconstruct the history of the effective population size from genomic data^14,53^, but in our system, changes in *N*_*e*_ predominantly arose from changes in the descendant number variance. To the extent that *σ*^2^ evolution occurs in other populations, inferred changes in *N*_*e*_ over time should not be assumed to reflect changes in census size alone. This may relate to the long-standing observation that *N*_*e*_ can have little relationship to census population size^16,20–22^, though we caution that other factors–particularly genetic draft from linked selection^**?**,54^–likely also contribute to that decoupling and may dominate in many natural populations.

The cause of the changes in descendant number variance we observe remains unclear, and several factors could in principle contribute. Technical variation in daily transfers could manifest as apparent excess 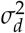, but such variation would also inflate differences between biological replicates. Because our replicate-to-replicate variation in estimated 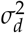 is manageably small (Figure 3C), technical variation does not appear to dominate the systematic trends we report. Spatial structure within the culture flasks could in principle inflate 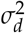, but the flasks are vigorously shaken throughout, so the populations are almost certainly well-mixed. A more likely source of excess descendant number variance is stochasticity in the growth process itself, including phenotypic heterogeneity among genetically identical cells.

In summary, tracking the strength of genetic drift across 50,000 generations of two LTEE populations revealed that the descendant number variance is not constant over evolutionary time in this system, with direct consequences for evolutionary dynamics. Together with prior work on sweepstakes reproduction, viral transmission, and microbial growth-trait variability, our findings reinforce the view that the descendant-number variance is a biologically important parameter that varies across populations and, at least in some cases, evolves within them. Whether *σ*^2^ evolution is similarly pronounced in other experimental systems and in natural populations remains to be tested. We hope this work motivates further investigation–both empirical and theoretical–into the causes and consequences of evolutionary changes in the strength of genetic drift.

## Methods

### Construction of Tn7 barcoded LTEE-derived clones

To make barcoded libraries for each LTEE-derived clone, we first determined the optimal agar plating media for each one that we wanted to barcode. We tested 1% agar plates made with either DM2000 (Davis Minimal Media supplemented with 2000mg/L glucose) or LB (Lysogeny Broth). Davis Minimal Media base (DM) consists of 5.36g/L potassium phosphate (dibasic), 2g/L potassium phosphate (monobasic), 1g/L ammonium sulfate, 0.5g/L sodium citrate, 0.01% Magnesium sulfate, 0.0002% Thiamine HCl. We chose the plating media that supported the most robust growth (Table S1), while also avoiding using LB for*S*and*L*clones, which we previously found to cause issues^55^. We generally used previously isolated and sequenced LTEE clones from Tenaillon et al. (2016)^29^, except for the*S*and*L*clones from 6.5k, 10k, and 50k generations. We used the 6.5k clones isolated from a different set of experiments^27^. For the 10k and 50k clones, we streaked whole population aliquots (REL4537 and REL11319, respectively) directly from glycerol stocks onto DM2000 plates. We then isolated several single colonies, resuspended them in DM2000 media, and used Sanger sequencing to sequence the*arcA*and*aspS*loci (Table S3), which each have a single nucleotide polymorphism that unambiguously distinguishes the*S*and*L*lineages. After confirming the lineage identity of the picked clones, we moved forward with a single*S*and*L*clone for each of the two populations.

We used a previously developed method^31^ to insert random DNA barcodes at the*attTn7*locus in the*E. coli* genome using a Tn7 transposon system. We used an older version of the Tn7 barcoding system that had a duplicated GFP-kanamycin resistance cassette. To make the barcoded Tn7 libraries for each LTEE-derived clone, we first streaked the LTEE strain for single colonies on the appropriate agar media, directly from glycerol stock. We also streaked the Tn7 “helper” donor strain (AMD1663) on LB/agar supplemented with 100µg/mL Carbenicillin and 0.3mM Diaminopimelic acid (DAP). The AMD1663 strain is an*E. coli*WM3064 donor with a pTNS2 plasmid (Addgene 64968), which encodes the Tn7 transposase genes^56^. We then picked single colonies for the LTEE clone and AMD1663, and grew them overnight in DM2000 and LB + 100µg/mL Carbenicillin + 0.3mM DAP, respectively. Additionally, we thawed an entire 1mL glycerol stock aliquot of AMD1386, a WM3064 donor containing the barcoded Kan-Tn7 transposon plasmid library, and grew overnight in LB + 100µg/mL Carbenicillin + 0.3mM DAP + 50µg/mL Kanamycin. The next day, we washed the two donor cultures (to eliminate the antibiotics) by centrifugation for 3 minutes at 2500 xg, aspirating the supernatant, and resuspending in DM0 (Davis Minimal Media without a carbon source) three times. We then combined and plated approximately 1 OD·mL of the LTEE culture, AMD1663, and AMD1386 on either LB + 0.3mM DAP plates or DM2000 + 50mM casamino acids (ThermoFisher; cat. no. 223050) + 0.3mM DAP plates, as appropriate. We grew the agar plates overnight to allow the two donor strains to conjugate with the LTEE clones, and initiate transposition of the Tn7 barcoded transposon. The following day, we scraped up the lawn on the agar plates into approximately 1.5mL of DM0, resuspended, and washed the culture by centrifugation for 3 minutes at 2500xg, aspirating the supernatant, and resuspending in DM0 three times (to eliminate residual DAP). We added sterile glycerol to a final concentration of 20%, and then serially diluted the mixture and plated on DM2000/LB + 50µg/mL Kanamycin agar plates, growing the plates overnight at 37^°^C. We saved the mixtures as glycerol stocks in the -80^°^C freezer. The following day, we determined the approximate total number of successful transconjugants by counting colonies. We then thawed the glycerol stock from the day prior, diluted the stock in DM0 so that we would expect to get approximately 10,000 colonies (and thus uniquely barcoded transconjugants), and plated the diluted mixture on DM2000/LB + 50µg/mL Kanamycin agar plates, growing the plates overnight at 37^°^C. The following day, we scraped up the colonies on the plate, resuspended in DM0 + 20% glycerol, and distributed the resuspended final library into four 1mL glycerol stock aliquots, saving them in the -80^°^C freezer.

### Genomic sequencing

The majority of genomes from LTEE clones used in this work have previously been sequenced–REL606^57^, the majority of both Ara-2 and Ara+2 clones^29^, and the 6.5k*S*and*L*clones ^27^. That left four clones that have not yet been sequenced–both the*S*and*L*clones from 10k and 50k generations. To sequence these clones, we grew them directly from glycerol stock in DM2000 media overnight. Then, we pelleted 2mL of the culture, and extracted genomic DNA with the DNeasy Blood and Tissue Kit (Qiagen 69504), according to the manufacturer’s protocol. Samples were prepared for whole genome sequencing using the Illumina DNA Prep tagmentation kit (cat. no 20018704) and IDT For Illumina Unique Dual Indexes (cat. no. 20027213-16). Sequencing occurred on the Illumina NextSeq2000 system using a 300 cycle flow cell kit, generating 2x150bp paired-end reads. To enhance base calling accuracy, 1-2% PhiX control was added to the sequencing run. Read demultiplexing, trimming, and analytical operations were conducted using DRAGEN v3.10.12, the integrated analysis software on the NextSeq2000. Reads were mapped to the REL606 genome^57^ and mutations called with `breseq`(v0.38.3) ^58^, all with default settings.

### BarSeq experiments

We performed two independent biological replicates of the BarSeq experiments for most strain libraries, done on different sets of days. For REL606, we performed three biological replicates. To start a BarSeq experiment, we first unfroze 1mL glycerol stock of the relevant transposon libraries and transferred the entirety to 10mL DM2000 media (supplemented with 50µL/mL Kanamycin) in a 50mL glass erlenmeyer flasks, which were grown for 16-24hrs at 37^°^C, shaken at 120rpm. Biological timecourse replicates using the same strain libraries were always performed on different sets of days (Table S1). All cultures for all experiments were grown with the same shaker, in the same 37C warm room. The next day, we washed the cultures by pelleting via centrifugation for 3 minutes at 2500 xg, aspirating the supernatant, and resuspending in DM0 three times. After thoroughly vortexing the cultures, we transferred them 1:1000 to 20mL DM25 (Davis Minimal Media supplemented with 25mg/L glucose) in a 50mL glass erlenmeyer flask. Subsequently, every 24 hours, we transferred 200µL of culture into 20mL of fresh DM25 in a 50mL glass erlenmeyer flask. Again, the cultures we grown at 37^°^C, shaken at 120rpm. We did this for three cycles to ensure that the cultures have sufficiently physiologically adapted to the culture conditions. Then, we repeated the procedure for five more days (Days 0-4), where we collected samples for later sequencing and colony counting. Pellets were harvested every day by centrifugation at approximately 20,000xg for 10min, using all of the culture remaining after transferring. Subsequently, the pellets were stored at -20^°^C until the experiment was finished. To measure population size for each culture at each time point via colony forming units (CFUs), we generally diluted cultures twice at 1:100 in 1mL of DM0, then we plated 100µL of the final diluted culture on an agar plate. This process was repeated so that each culture had two independent “technical” plating replicates. We used either LB or DM2000 agar plates, again depending on which media allowed the strain in question to form the most robust colonies (Table S1).

After the completion of the experiment, pellets were retrieved from the -20^°^C freezer, and genomic DNA was extracted using the Qiagen DNeasy blood and tissue extraction kit (cat no. 69504), with elution in double-distilled water yielding approximately 50ng/µL of DNA. DNA Tn7 barcodes were amplified from genomic DNA samples using PCR with Q5 Hot Start Polymerase (NEB, cat. no. M0493S); the 50µL reactions included 5µL PCR primers, 5µL genomic DNA, 10µL 5x buffer, 10µL GC enhancer, 1µL dNTPs, 0.5µL Q5 polymerase, and 18.5µL water. Custom dual-indexed primers were utilized, incorporating binding sites adjacent to the barcode region, along with required Illumina read/index binding sites^59,60^. The forward primer sequence was AATGAT ACGGCG ACCACC GAGATC TACACT CTTTCC CTACAC GACGCT CTTCCG ATCT N_*n*_XXXXXX GTCGAC CTGCAG CGTACG, where X represents the custom forward 6bp index, and N_*n*_ indicates 1-4 random nucleotides, varying by primer pair; the reverse primer sequence was CAAGCA GAAGAC GGCATA CGAGAT XXXXXX GTGACT GGAGTT CAGACG TGTGCT CTTCCG ATCTGA TGTCCA CGAGGT CTCT, where X denotes the standard Illumina 6bp IT index. A distinct primer pair was used for each genomic DNA sample from different experiments, replicates, or time points to enable demultiplexing after sequencing. The PCR protocol involved 4 minutes at 95°C, followed by 25 cycles of [30 seconds at 95°C, 30 seconds at 55°C, 30 seconds at 72°C], and a final extension of 5 minutes at 72°C. The correct PCR products were confirmed through agar gel electrophoresis. All PCR reactions were subsequently pooled and purified using the Zymo DNA Clean and Concentrator kit (cat. no. D4013), eluted in double distilled water. The final pooled sample was sequenced on an Illumina HiSeq 4000 (50SR) at the Vincent J. Coates Genomics Sequencing Laboratory at UC Berkeley.

### Barcode sequencing data preprocessing

To process the raw Illumina sequence reads for each sample, we first employed a previously described Perl script (`MultiCodes.pl`)^59,60^ to process the demultiplexed sequencing reads. This script extracts the barcode sequences by trimming the areas associated with the sequencing primers and the regions flanking the barcode. It also excludes reads that fail to align with the secondary sequencing index or that do not meet the requisite quality scores. Subsequently, we tally the counts of unique barcodes to compile a table linking each barcode sequence with its corresponding counts.

However, errors during PCR and sequencing can lead to mutations in some barcode reads, which can be incorrectly interpreted as distinct and real barcodes. Therefore, it is necessary to correct these sequencing errors by merging mutated barcodes with their original versions and consolidating their read counts. We do this by creating a graph where each node is a barcode, drawing an edge if the Levenshtein (edit) distance between the two barcodes is ≤ 3; to populate the nodes of the graph, we use the union of all barcodes from each time point in an experiment. We excluded barcodes with highly skewed GC contents–those lower than 20% or higher than 80%. The previously mentioned Perl script detects barcode pairs that differ by one mutation; we initialize the graph by drawing edges between these barcode pairs. For all remaining pairs of barcodes, to avoid excessively long run times, we first computed their Hamming distance. We then only computed the Levenshtein distance between a barcode pair if the Hamming distance ≤ 10. We then populate the edges of the barcode similarity graph using networkx^61^. Finally, we pull out all connected components of the graph, and treat each connected component as a “superbarcode,” summing together the read counts for each time point from each constituent barcode. For all subsequent analyses, we proceed to use the “superbarcodes” read trajectories, and will generally refer to them simply as “barcodes.”

### Outlier detection and barcode filtering

We observed some barcodes appear to have non-neutral fitness effects, likely due to spontaneously occurring selected mutations on the background of the barcoded lineage. We thus sought to detect and filter out barcodes with putatively non-neutral fitness effects, using a simple likelihood-based model. Using an approach similar to that presented in Ba et al. (2019)^62^, we model sequencing read trajectories for barcode 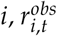, as a negative binomial random variable parameterized such that the variance is proportional to the mean,

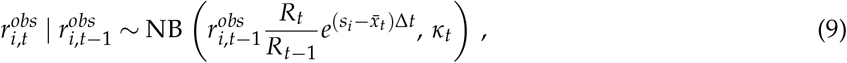

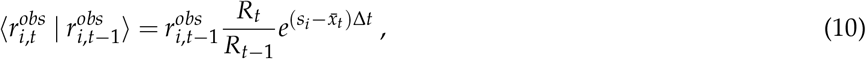

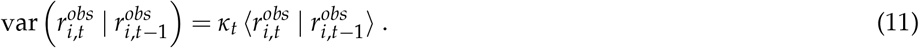

We chose to use a negative binomial density to more accurately capture the discreteness of sequencing reads, which is important for more accurately modeling barcodes with low read counts. Here, *R*_*t*_ is the total number of sequencing reads (across all barcodes) at time point *t, κ*_*t*_ is total over dispersion arising from a combination of sequencing measurement error and genetic drift, *s*_*i*_ is the fitness effect of barcode *i*, 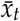 is the average fitness of the population at time *t*, and Δ*t*=6.64 is the number of generations per cycle. Before inferring the fitness effect for each barcode, we estimate the overdispersion parameters *κ*_*t*_ and the population-averaged fitness 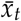 from all barcodes in the experiment. We first roughly filter the barcodes so that their frequency is (i) large enough so that the central limit theorem applies to both read counts and the number of cells per barcode at the bottleneck, and (ii) is small enough that they are less likely to have a fitness effect (larger lineages appear more likely to be non-neutral). Thus, for inference of *κ*_*t*_ and 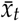 we chose to focus on barcodes with observed frequencies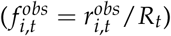at time *t*− 1above max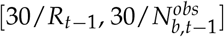 and below7 · 10^−4^. Here, 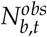 is the bottleneck size estimated from the CFU data (averaged between the two technical replicates), 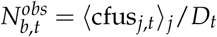. The inferred overdispersion and mean fitness parameters are not sensitive to changes in the filtering cutoffs, and we use robust statistics to infer the two parameters. Using the remaining barcodes after filtering, we estimated the mean fitness for each timepoint as,

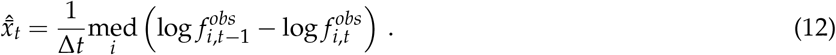

We again used a robust estimation method for *κ*_*t*_^33^. First, we used a square root variance-stabilizing transformation on the observed frequencies to eliminate the mean-variance dependence, and took the difference between the two neighboring time points, 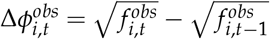. We use the median absolute deviation to robustly estimate the variance and then scale appropriately to recover*κ* _*t*_,

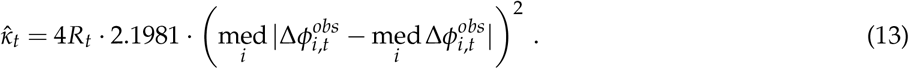

We used standard bootstrapping to estimate the sampling distribution of both parameters. The relative parameter standard errors were generally ≲ 1%, so we proceeded with the parameter point estimates. We note that we only use these two inferred parameters for the purpose of outlier detection, and not for any other downstream application.

We can now proceed to infer fitness effects independently for each barcode via maximum likelihood. In order to capture both constant fitness effects and time-varying fitness effects (arising due to e.g. fluctuating selection, or multiple different selected lineages within a barcode), we obtain maximum likelihood estimates of *s*_*i*_ when all timepoints are combined,

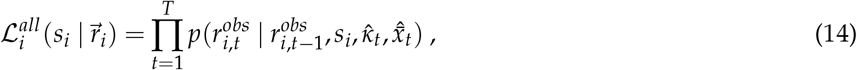

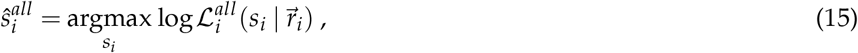

along with the estimate for each time point pair,

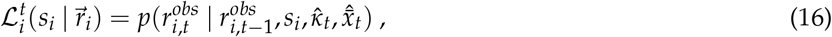

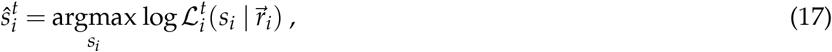

in both cases using the density from equation 9. We then compute a p-value for the fitness effect of each barcode under the null hypothesis *s*_*i*_ = 0, for all estimates, 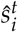 and 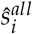 . For ease of computation and generality we compute the p-value as the posterior probability that the likelihood ratio between null and alternative hypotheses is greater than 1, i.e. the probability that the data more strongly support the null hypothesis over the alternative^33^,

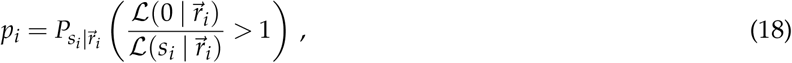

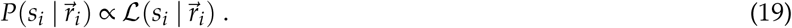

This convenient definition has been shown to be equivalent to the frequentist definition of the p-value using a likelihood ratio test statistic (if the distribution is invariant under transformation)^63,64^, and does not require asymptotic approximations. We computed separate p-values using 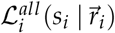 and all 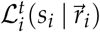 for the likelihoods.

In practice, this p-value can be calculated by first, finely discretizing the likelihood curve along *s*_*i*_ and normalizing it to get the posterior,

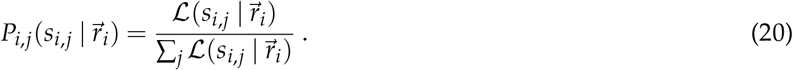

Then, calculating the log-likelihood ratio along all discretized *s* values,

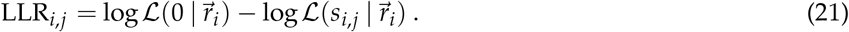

And finally, summing to get the posterior probability that the data supports the null hypothesis more than the alternative, where*I*[·]is the indicator function,

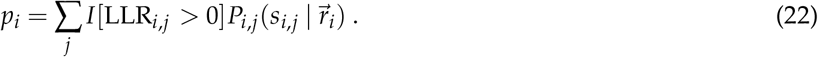

We then want to exclude barcodes that have likely non-neutral fitness effects from further analysis. We first performed an initial (liberal) barcode filtering where we filter all barcodes where the observed frequency at day 0 is above the 50th percentile, and below the 10th percentile; we did this because we had observed that high-frequency barcodes are more likely to have apparent non-neutral fitness effects, and low-frequency barcodes are more likely to be sequencing artifacts. We then choose to filter out barcode *i* if it had any p-values (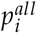and 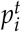 for all *t*) less than 0.05. We chose this p-value cutoff (*α*) because it seems to represent the near-optimal balance between true and false positive rates in simulations of the outlier detection and genetic drift inference pipeline (Figure sS10-S9). We are not using the inferred p-values to claim that any barcodes that fall under the cutoff are under selection with any significant certainty; we are simply trying to exclude as many putatively selected barcodes as possible, while minimizing the exclusion of truly neutral barcodes. On average, this filtering excluded approximately 15% of barcodes per library from further analysis (Table S2). We find that our results are robust to our choice in p-value cutoff (Figure S8).

Following filtering, we sum the remaining low-frequency barcodes together so that each (combined) barcode is always at sufficiently high frequency such that the central limit theorem applies to the trajectory statistics. We do this by first computing a time-dependent cutoff frequency,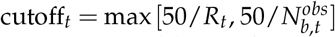. For each time point, we sum the lowest barcode below the cutoff with the trajectory of the second-lowest barcode. We iteratively continue this process until no barcodes remain under any of the cutoffs at any time.

### Estimates of mean squared displacements

We expect that the mean squared displacement (MSD) of the square-root transformed barcode frequency trajectories (Figure 1F) should increase linearly over time increments with a slope determined by *N*_*e*_ (assuming that *N*_*e*_ does not change over time),

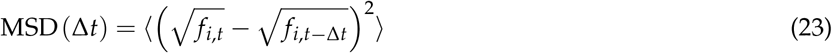

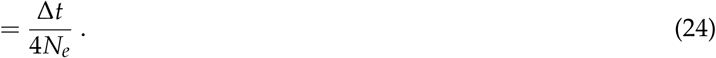

We estimated the MSD for all pairs of time points that were either 1, 2, or 3 time increments (Δ*t*) apart, using barcodes that passed filtering (from the prior section). We require a method to distinguish between measurement error (which is uncorrelated in time) and genetic drift (which accumulates over time). We applied a method previously employed^30,33^ to differentiate between these two types of noise. In brief, the variance between two time points represents the sum of the decoupling noise and the measurement noise (*ζ*_*t*_) at those time points,

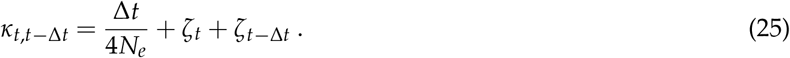

For each barcode, we calculate the difference of square-root transformed frequencies between considered pairs of time points. We then obtain a robust estimate of 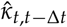 by using the median absolute deviation (equation 13), across all barcodes. We estimate the standard error, std, 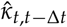, through standard bootstrapping techniques. Using the relationship outlined in equation 25 between *κ*_*t,t*−Δ*t*_ and the noise parameters, we can determine the noise parameters based on our measured values of 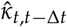 . This is achieved by numerically minimizing the weighted squared differences, expressed as 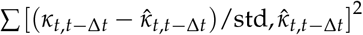. Our final estimate of the MSD is obtained by adjusting each 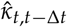 for the measurement noise parameters, and then averaging the adjusted values across all instances with the same time increment Δ*t*,

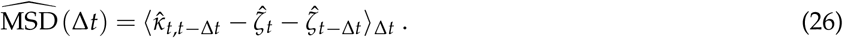

We obtain MSD confidence intervals via bootstrapping.

### Bayesian inference of genetic drift parameters

Following Yu et al. (2024)^34^, we model the frequency trajectories of (post-filtering) lineages within a population using a Cannings-type model. If the number of individuals within lineage *i* is sufficiently large such that the central limit theorem applies, and each individual within the lineage behaves identically and independently, we can model the lineage frequency dynamics,*f* _*i,t*_, as the following Markov chain,

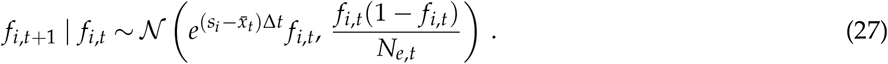

Here, *s*_*i*_ is the relative fitness effect of barcode *i*, 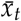 is the average fitness of the population at time *t*, Δ*t*≈6.64is the number of generations per cycle, and *N*_*e,t*_ is the effective population size at time *t*. The effective population size can be decomposed into two components–the bottleneck population size, *N*_*b*_, and the descendant number variance, 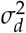. We assume throughout that the bottleneck population size can vary over time, but the descendant number variance is constant, i.e. 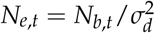 . We further simplify the model by restricting ourselves to small lineage frequencies, *f*_*i,t*_ ≪ 1, such that we can ignore the1 −*f* _*i,t*_ nonlinearity, and only considering neutral lineages, *s*_*i*_ = 0,

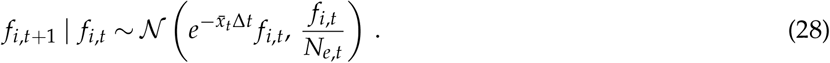

In our experiments, we never directly observe the frequency of a lineage; we indirectly measure the frequency by sequencing the barcode regions and counting the barcodes. Thus, the measurement process will introduce additional noise that will be at least as strong as Poisson sampling noise. If (i) the observed number of sequencing reads, 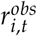, is sufficiently large such that the central limit theorem applies, (ii) the measurement process is unbiased, and (iii) measurement noise affects every barcoded cell/sequencing read independently and identically, we can model measurement as the following,

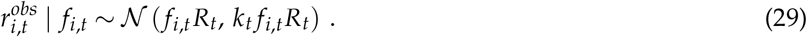

Here, *R*_*t*_ is the total number of sequencing reads (across all lineages) at timepoint *t*, and *k*_*t*_ is sequencing measurement error overdispersion. This mean-variance relationship was confirmed by analyzing the noise in BarSeq technical replicates from a previous experiment^33^ with the exact same sample processing procedure that we use here (Figure S6). The measurement error is solely driven by poisson sampling noise when *k*_*t*_ = 1; any other sources of noise (e.g. associated with gDNA extraction or the amplicon PCRs) will increase *k*_*t*_ above one. We can rewrite equation 29in terms of the observed frequency,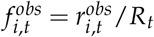,

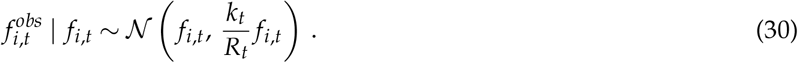

Together, equations 28 and 30 define a hidden Markov model, which can be used to infer parameters from collected data. To ease further analysis, we apply a square-root variance stabilizing transformation on both equations, where we define 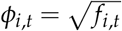,

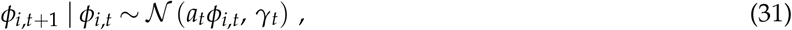

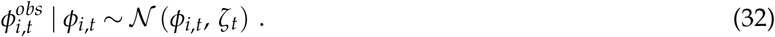

We expect this variance stabilizing transformation to be valid as long as*ϕ* _*i,t*_ is sufficiently large. For notational simplicity, we defined the following compound parameters,

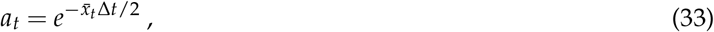

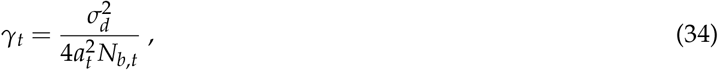

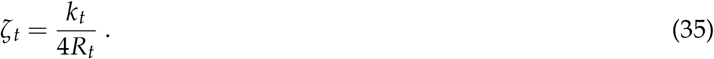

With the mean-variance dependency extinguished, we can treat the system as a standard Kalman filter, integrating out the unobserved*ϕ* _*i,t*_ states, and obtaining a recursively-defined likelihood,

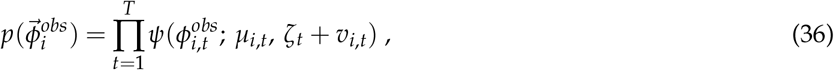

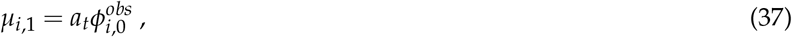

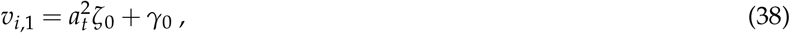

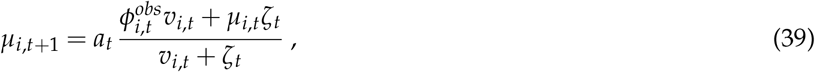

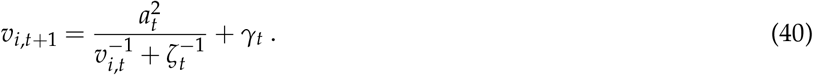

We assumed that we obtained measurements of 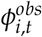at every time point from time 0 to *T*. We defined *ψ*(*x* ; *µ, v*) as a Gaussian density with mean *µ* and variance *v*.

In order to separate the contribution of *N*_*b,t*_ and 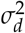 to *N*_*e,t*_, we would need measurements of how the bottleneck population size *N*_*b,t*_ changes over time. To that end, every day in the timecourses from days 0 to 3 we plated the end-of-cycle cultures on agar plates at a defined dilution rate, with two technical replicates for each culture. Once colonies grew, we counted them and used the colony forming units (CFUs) as a measure of the bottleneck population size. We then modeled the CFUs as a negative binomial random variable, parameterized such that the variance is proportional to the mean,

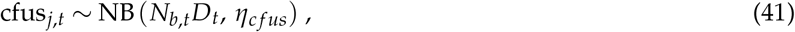

Here *j* ∈ {1, 2} represents the technical replicate. Here *D*_*t*_ is the total dilution rate and *η*_*c f us*_ is the measurement error overdispersion. Again, when *η*_*c f us*_ = 1, the noise reduces to poisson sampling error. We did not obtain a technical replicate for certain time points/cultures (due to e.g. plate contamination), in which case we only used the CFU measurement from the single replicate. We experimentally verified the proposed mean-variance relationship for CFUs, and obtained a preliminary estimate of *η*_*c f us*_; we grew three independent cultures of REL606 at different cell densities, then we diluted and plated 16 independent technical replicates for each culture. We used the same dilution and plating protocol as the main experiments. We then computed the CFU mean and variance across technical replicates for each culture. We observed a linear mean-variance relationship for the CFUs, as expected, with a slope greater than one (Figure S5A). We sought to incorporate this experimental data into our model. Thus, we used the Hamiltonian Monte Carlo (HMC) algorithm implemented in STAN^65^ to jointly estimate the posterior of each parameter in the following model,

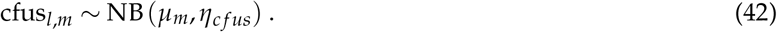

Here, *l* is the label for each CFU measurement, *m* ∈ {1, 2, 3} is the label for each independent culture at a different density, and *µ*_*m*_ is the underlying “true” culture density for each group. We used a uniform prior for each *µ*_*m*_ and a Pareto prior for the overdispersion parameter 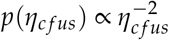 . We found that the marginal posterior of *η*_*c f us*_ was well fit by a gamma distribution. We thus fit the posterior estimate to a gamma distribution via maximum likelihood, and obtained estimates of the gamma distribution parameters (Figure S5B),

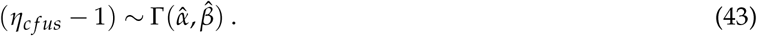

We can now use equations36-41as the likelihood for our model describing lineage frequency trajectories in our timecourse experiments. We combine the likelihoods from all barcodes in the experiment, and raise each likelihood to the power of *ξ* (“power posterior”), 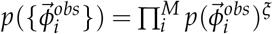. We use the power posterior on the suggestion of Miller and Dunson (2019)^66^ as a source of mild regularization, increasing the relative weight of the priors. Here, *ξ*= [1 +*M*/*α*] ^−1^, where we chose *α*= 500, as simulations indicate that it is the largest (i.e. most conservative) value that still has a (small) regularizing effect. To perform inference on the model, we again use Pareto priors for the relevant overdispersion parameters of the model, 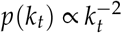 and 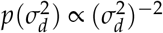. We use a weakly regularizing, prior for 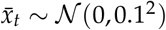 as relative fitness effects generally do not exceed 0.1/generation in LTEE strains and environment^*33,67*^. We use equation 43 as the prior for *η*_*c f us*_ (with the maximum likelihood-fit parameters), allowing us to incorporate the information from the CFU technical replicates experiment, while also allowing the posterior of *η*_*c f us*_ to change based on the data for each timecourse. And we use uniform priors for all *N*_*b,t*_. We then used the priors and likelihoods along with the HMC algorithm implemented in STAN to jointly estimate the posterior of each parameter. We used the same inference procedure to obtain final parameter estimates, where the data from the biological replicates is pooled together; all parameters except 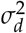 and *η*_*c f us*_ were allowed to vary independently for data from each replicate. All HMC chains were evaluated for convergence by ensuring that the 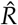 statistic was between 1 and 1.01, and through visual inspection.

We performed all of our BarSeq experiments in a culture volume two times larger than the LTEE cultures (20mL versus 10mL), but retaining the same dilution rate and all other culture conditions. This was primarily to be able to obtain a sufficiently large sample for genomic DNA extraction. We thus divided our estimates of effective population size and bottleneck population size by a factor of two, to enable more direct comparison with the standard LTEE culture environment.

### Simulations of the data generating process and inference pipeline

We ran simulations of the data generating process to validate our inference pipeline and probe the effects of the regularization measures. We directly modeled how the number of cells *n*_*t*_ in a lineage changes over time, where the markov transition density is a negative binomial with the variance proportional to the mean,

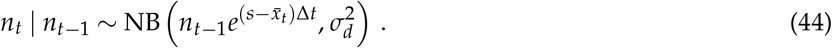

We always initialize the number of cells with *n*_0_ = 500, and use 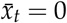and Δ*t*= 6.64generations. We simulate the number of sequencing reads as a negative binomial random variable, again where the variance is proportional to the mean,

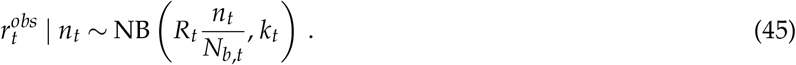

The observed frequencies are then 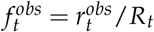. In our simulations, we always use *N*_*b,t*_ = 5 · 10^6^, *R*_*t*_ = 5 · 10^6^, 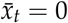, and *k*_*t*_ = 2, which are typical parameters for our experimental data. We varied the descendant number variance, 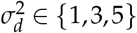. For each simulated dataset, we independently simulated 1000 barcodes. Each barcode has a 20% chance of having a non-neutral (non-zero) fitness effect; this percentage of selected barcodes is likely high relative to our experimental data, but we we interested in exploring how robust our pipeline is, even in the presence of a large amount of “contaminating” data. If the barcode is randomly selected to have a non-zero fitness effect, there is a 50% chance it will have a negative (deleterious) fitness effect, and a 50% chance it will have positive (beneficial) fitness effect. Beneficial and deleterious fitness effects, respectively, are drawn from the following two exponential distributions,

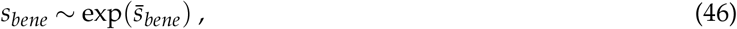

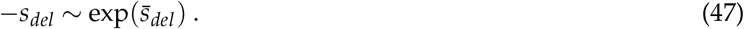

We chose exponential distributions because previously measured distributions of fitness effects (DFEs) for LTEE strains can be well approximated by exponential distributions^33^. Ascensao et al. (2023) found that the deleterious DFEs of REL606, 6.5k*S*and 6.5k*L*in monoculture (the same condition studied in this work) were all approximately this same distribution, with 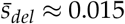. By contrast, the beneficial DFE in REL606 was higher, with 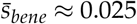, and the beneficial DFE of 6.5k*S*and*L*lower, with 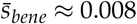. Thus, for all simulations, to accurately reflect the spectrum of mutations that the LTEE populations likely have available, we always used 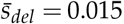, and used either 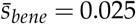 or 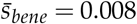.

For each simulation run, we simulated CFUs as a negative binomial with the variance proportional to the mean, with two “technical replicates” per time point,

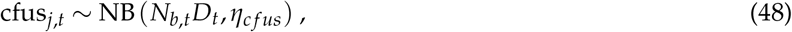

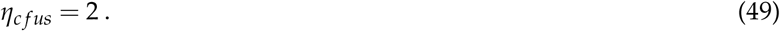

To summarize, we simulated 100 independent “experiments” for each of six distinct parameter sets, where we varied 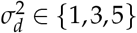 and 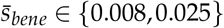. For each experiment, we obtained simulated CFU data. Within each experiment, we simulated 1000 independent barcode sequencing read trajectories, 20% of which (on average) have non-neutral fitness effects.

Once we obtained all simulated datasets of observed frequency trajectories and CFUs, we pushed the data through the fitness inference/outlier detection pipeline and filtered the barcodes appropriately, as in the real experiments. The outlier detection method appears to be able to detect non-neutral lineages sufficiently well; we chose a p-value cutoff of 0.05 that balances a low false positive rate with a high enough true positive rate (Figure S9). Then we used the previously introduced Bayesian inference method to jointly infer the genetic drift parameters, i.e. *N*_*b,t*_ and 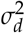, along with all the nuisance parameters, i.e. *k*_*t*_, *η*_*c f us*_, and 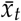. Additionally, we ran the Bayesian inference in the case where we did not use either or both of the robustifying methods, i.e. outlier detection or the regularizing power posterior (Figure sS10-S11).

### Simulations of noisy growth processes

We simulate noisy asexual splitting growth processes to compare to our analytical results (Figure 5A-B; SI section S3). To simulate variable lag times, we sampled about10 ^7^ independent instances of *t*_*lag*_ from either a gamma (Figure 5B) or exponential (Figure 5B) distribution, for a given set of parameters. After the lag time, each individual starts growing exponentially (deterministically), such that 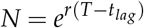. We use *T*= 12hrs and *r*= 0.03min ^−1^, which is a typical growth rate for*E. coli* ^41,42^. Thereafter, we compute the descendant number variance by appropriately averaging over the simulated dataset,

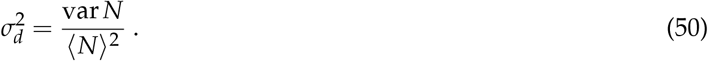

To simulate noise in doubling times, we simulated10 ^5^ independent lineages, each starting from a single individual. We draw the doubling time from either a gamma (Figure 5A) or exponential (Figure 5A) distribution. At doubling time, the individual splits into two new individuals, and new random doubling times are drawn for each of the new individuals. This process is continued until time *T*, which we take as 6.64 generations. The simulated distribution of the total number of individuals per lineage is then used to compute the descendant number variance, by again applying equation 50.

### Evolutionary dynamics simulations

We simulated the dynamics of the first mutational step, when mutations can be drawn from a joint distribution of fitness effects and descendant number variances, 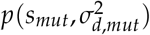 (Figure 5C-F). In the main text, we took the joint distribution to be a Gaussian copula with exponential marginals **??**. Let (*Z*_1_, *Z*_2_) be standard bivariate normal with correlation *ρ*∈[−1, 1]. Define

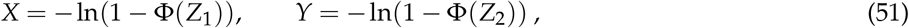

where Φ is the standard normal CDF. Then *X, Y* ∼ Exp(1) marginally, and as *ρ* ranges over [-1,1], the dependence between *X* and *Y* ranges from countermonotonic (*ρ* = -1, giving Pearson correlation *π*^2^/6) through independence (*ρ*=0) to comonotonic (*ρ*=1, giving Pearson correlation 1).

We also took the joint distribution to be Gumbel’s bivariate exponential distribution^68^ (Figure S16), where the density of the distribution is,

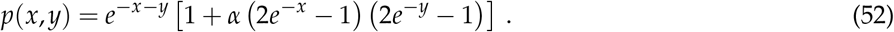

Under this joint distribution, both *x* and *y* have exponential marginal distributions with mean of one. Here, *α* is a parameter that controls the strength of correlation between *x* and *y*, which is directly related to Pearson’s correlation coefficient, *ρ*_*p*_ =*α*/ 4. We vary *α*∈ {− 1, 0, 1}. We also chose a joint distribution where *s*_*mut*_ and 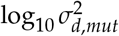are uncorrelated Gaussian distributions (Figure S17).

After sampling from a given joint distribution, we scale *x* and *y* to their desired means and locations. We chose a mean fitness effect of 0.025, as previous work has shown that the distribution of fitness effects of REL606 can be well approximated as an exponential distribution with mean around 0.025^33^.

To simulate the evolutionary dynamics in the strong selection, weak mutation regime (periodic sweeps), we can consider the fate of each mutation independently. To that end, for10 ^6^ independent replicates, we initialize a mutant at a population size of 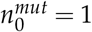 and the wildtype at 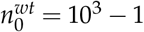. For each mutant, we sample a pair of *s*_*mut*_ and 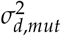 from the aforementioned joint distribution. For the wildtype, we always use *s*_*wt*_ = 0(by definition) and 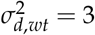. We then simulate the population using a Wright-Fisher like procedure, with an expected total population size of *N*_*tot*_ = 10^3^. Every generation, we compute the expected population size for both the mutant and wildtype, and renormalize to obtain an expected strain frequency,

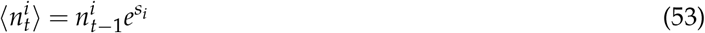

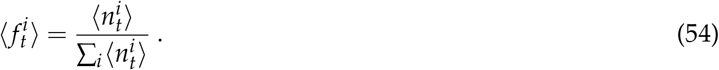

Subsequently, for each strain, we simulate the effect of genetic drift by sampling the population size of the next generation for each strain*i*from a negative binomial density (similar to equation 44),

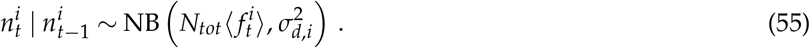

We iteratively continue this procedure until either the mutant or wildtype goes extinct. We then take all of the simulation runs where the mutant successfully fixed in the population, and average across the mutant parameter values to obtain ⟨*s* _*mut*_|fixed⟩ and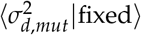.

We used a slightly different simulation procedure to simulate evolutionary dynamics in the weak selection, strong mutation regime (clonal interference). We initialize the population with solely the wildtype at *n*^*wt*^ = 10^3^. Then, for each generation, each individual has a probability of *U*_*b*_ = 10^−3^ of obtaining a beneficial mutation, drawn from the specified joint distribution distribution. We assume that fitness effects are additive, and the mutant descendant number variance replaces that of its ancestor. We then used equations53-55to simulate the dynamics of each strain across generations. We continue each simulation until one (non-wildtype) lineage has fixed in the population. We then take the root mutant parameter values of the lineage that eventually fixed, and average over all independent simulations to obtain ⟨*s* _*mut*_|fixed⟩ and 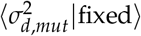.

## Code and data availability

Raw genomic sequencing reads are available at BioProject PRJNA1129587. All code and data presented in this manuscript are available at:https://github.com/joaoascensao/genetic-drift-evolution

## Acknowledgements

We thank Robin Herbert and Adam Deutschbauer for gifting us the barcoded Tn7 donor library, and providing advice on the system and protocols. We thank Adam Arkin, Kelly Wetmore, Aditya Prasad, Takashi Okada, and all members of the Hallatschek lab (past and present) for helpful comments and advice on the project. We thank Richard Lenski for sending us the LTEE-derived strains and populations, along with experimental advice and feedback. Research reported in this publication was supported by a National Science Foundation CAREER Award (1555330). This work was supported by the National Institute of General Medical Sciences of the NIH under award R01GM115851 and by a Humboldt Professorship of the Alexander von Humboldt Foundation. JAA acknowledges support from an NSF graduate research fellowship, a Berkeley fellowship (from UC Berkeley), and Lloyd and Brodie scholarships (from UC Berkeley Dept of Bioengineering). QY acknowledges support from an NSF graduate research fellowship. Genomic sequencing was performed by SeqCoast Genomics LLC. This work used the Illumina HiSeq 4000 at the QB3 Sequencing Laboratory at UC Berkeley (RRID:SCR_022170) for barcode sequencing, supported by NIH S10 OD018174 Instrumentation Grant.

## Supplementary Information

### S1 Supplemental Tables

**Table S1.**
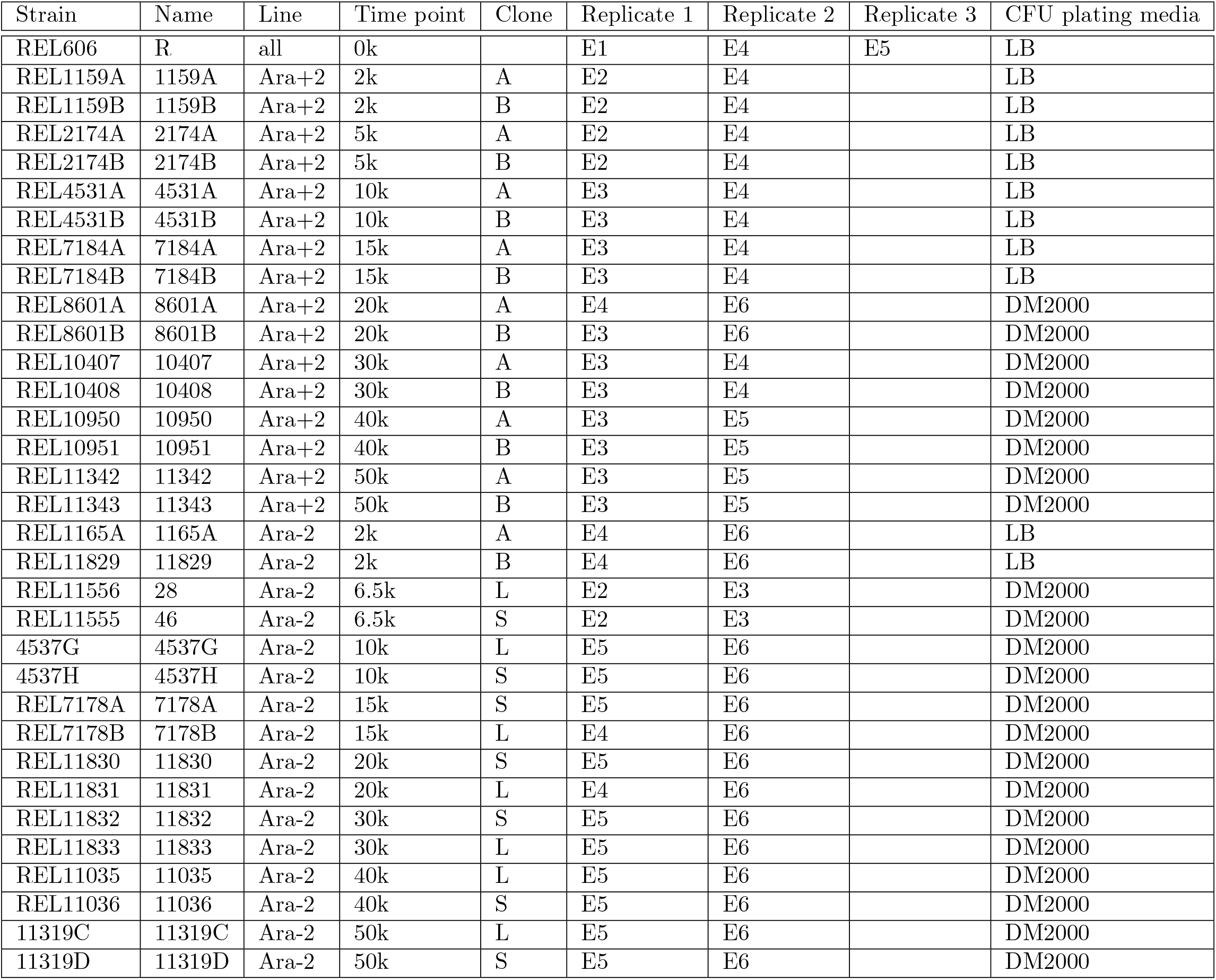
Table showing when replicates of different Tn7 barcoded timecourses were run. Each set of experiments E1-6 was run separately, on different sets of days.

| Strain | Name | Line | Time point | Clone | Replicate 1 | Replicate 2 | Replicate 3 | CFU plating media |
| --- | --- | --- | --- | --- | --- | --- | --- | --- |
| REL606 | R | all | 0k |  | E1 | E4 | E5 | LB |
| REL1159A | 1159A | Ara+2 | 2k | A | E2 | E4 |  | LB |
| REL1159B | 1159B | Ara+2 | 2k | B | E2 | E4 |  | LB |
| REL2174A | 2174A | Ara+2 | 5k | A | E2 | E4 |  | LB |
| REL2174B | 2174B | Ara+2 | 5k | B | E2 | E4 |  | LB |
| REL4531A | 4531A | Ara+2 | 10k | A | E3 | E4 |  | LB |
| REL4531B | 4531B | Ara+2 | 10k | B | E3 | E4 |  | LB |
| REL7184A | 7184A | Ara+2 | 15k | A | E3 | E4 |  | LB |
| REL7184B | 7184B | Ara+2 | 15k | B | E3 | E4 |  | LB |
| REL8601A | 8601A | Ara+2 | 20k | A | E4 | E6 |  | DM2000 |
| REL8601B | 8601B | Ara+2 | 20k | B | E3 | E6 |  | DM2000 |
| REL10407 | 10407 | Ara+2 | 30k | A | E3 | E4 |  | DM2000 |
| REL10408 | 10408 | Ara+2 | 30k | B | E3 | E4 |  | DM2000 |
| REL10950 | 10950 | Ara+2 | 40k | A | E3 | E5 |  | DM2000 |
| REL10951 | 10951 | Ara+2 | 40k | B | E3 | E5 |  | DM2000 |
| REL11342 | 11342 | Ara+2 | 50k | A | E3 | E5 |  | DM2000 |
| REL11343 | 11343 | Ara+2 | 50k | B | E3 | E5 |  | DM2000 |
| REL1165A | 1165A | Ara-2 | 2k | A | E4 | E6 |  | LB |
| REL11829 | 11829 | Ara-2 | 2k | B | E4 | E6 |  | LB |
| REL11556 | 28 | Ara-2 | 6.5k | L | E2 | E3 |  | DM2000 |
| REL11555 | 46 | Ara-2 | 6.5k | S | E2 | E3 |  | DM2000 |
| 4537G | 4537G | Ara-2 | 10k | L | E5 | E6 |  | DM2000 |
| 4537H | 4537H | Ara-2 | 10k | S | E5 | E6 |  | DM2000 |
| REL7178A | 7178A | Ara-2 | 15k | S | E5 | E6 |  | DM2000 |
| REL7178B | 7178B | Ara-2 | 15k | L | E4 | E6 |  | DM2000 |
| REL11830 | 11830 | Ara-2 | 20k | S | E5 | E6 |  | DM2000 |
| REL11831 | 11831 | Ara-2 | 20k | L | E4 | E6 |  | DM2000 |
| REL11832 | 11832 | Ara-2 | 30k | S | E5 | E6 |  | DM2000 |
| REL11833 | 11833 | Ara-2 | 30k | L | E5 | E6 |  | DM2000 |
| REL11035 | 11035 | Ara-2 | 40k | L | E5 | E6 |  | DM2000 |
| REL11036 | 11036 | Ara-2 | 40k | S | E5 | E6 |  | DM2000 |
| 11319C | 11319C | Ara-2 | 50k | L | E5 | E6 |  | DM2000 |
| 11319D | 11319D | Ara-2 | 50k | S | E5 | E6 |  | DM2000 |

**Table S2.**
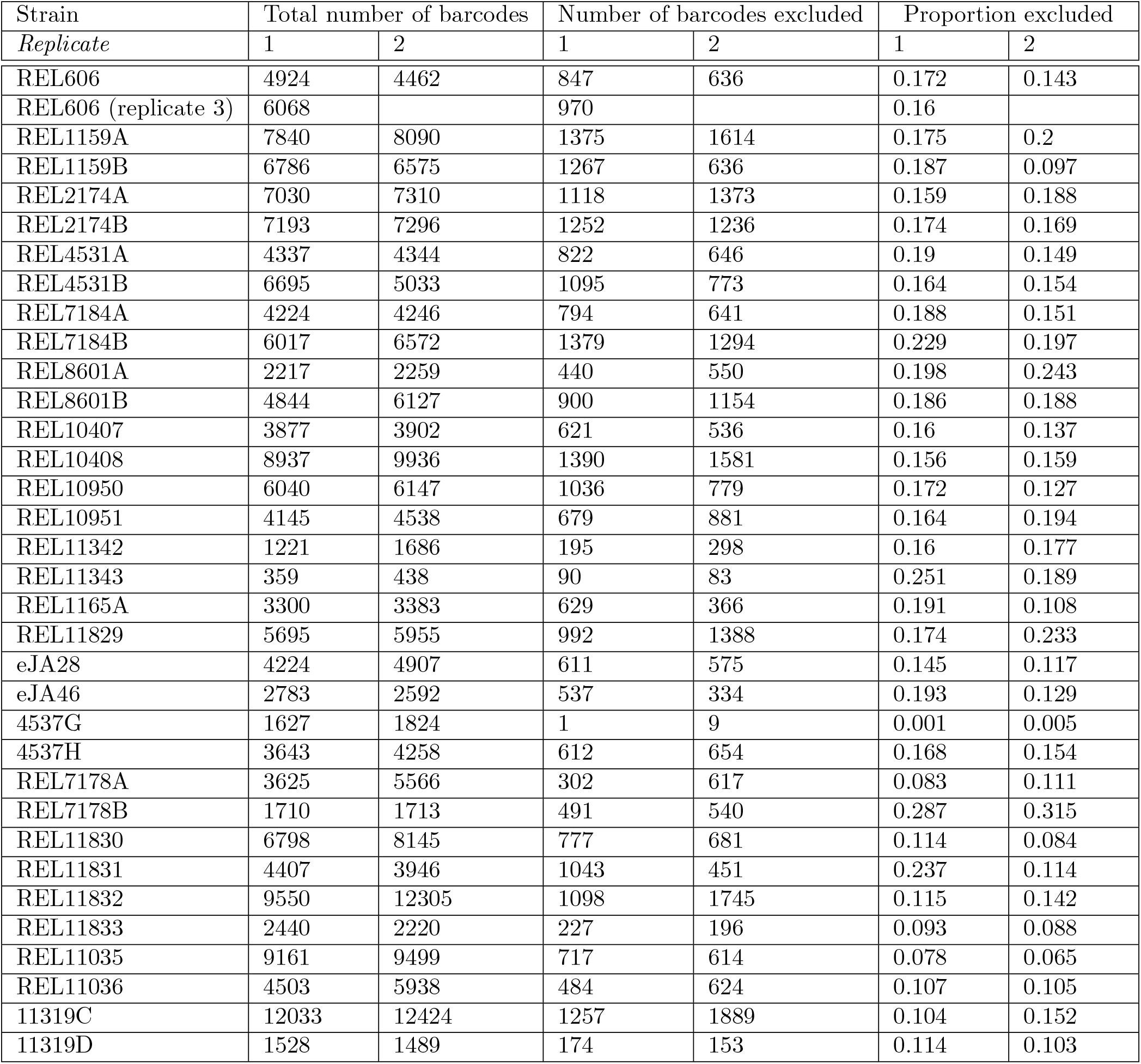
Summaries of excluded barcodes from likelihood-based outlier filtering (see Methods subsection “Outlier detection and barcode filtering”), where the detection threshold is*α*= 0.05. We show the total number of barcodes before outlier exclusion, the number of barcodes excluded, and the proportion of barcodes excluded for each tested library and biological replicate. We report the values for the 3rd replicate of the REL606 library as a separate row instead of adding new columns for compactness of representation.

**Table S3.**
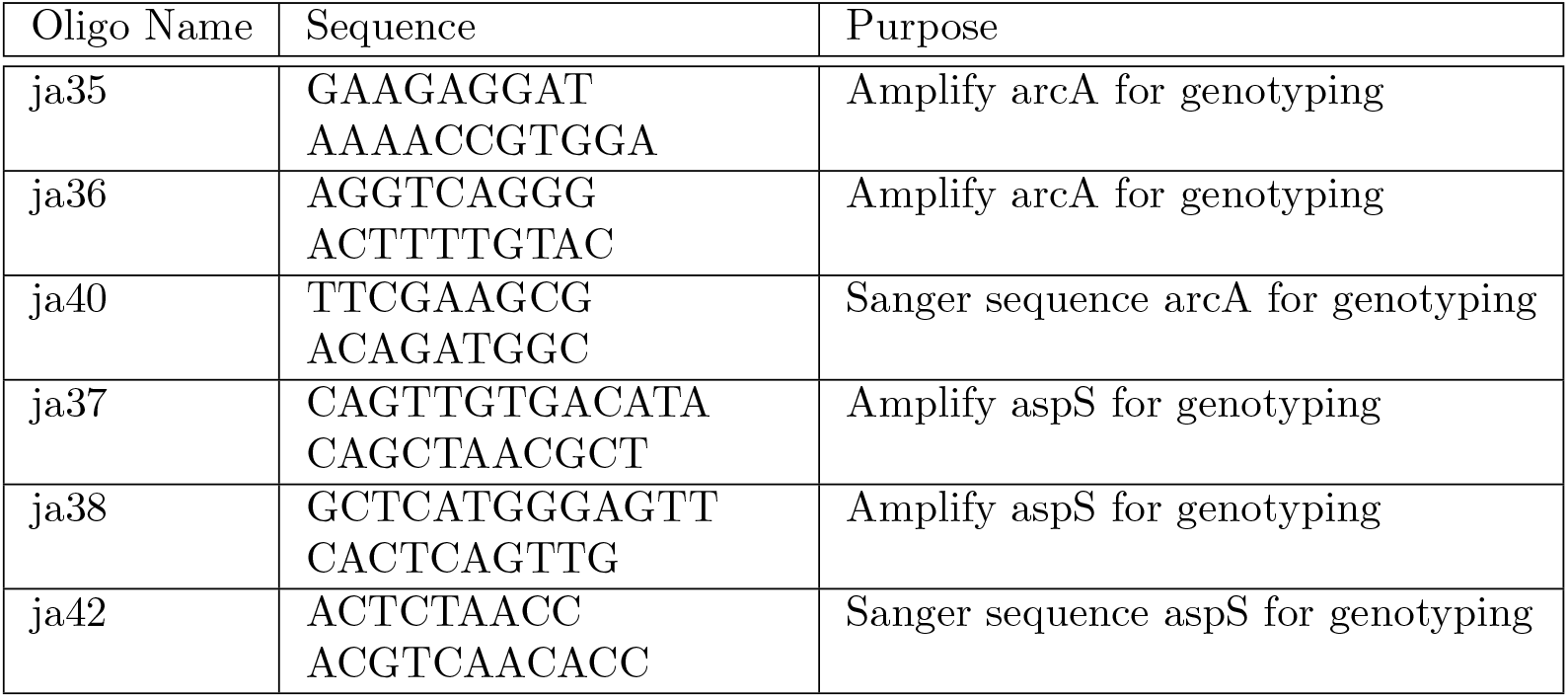
Table of genotyping oligonucleotides to distinguish*S*and*L*clones. [1].

### S2 Supplementary Note: Relation to Wahl and Gerrish (2001)

This note clarifies the relationship between the parameterization used in our manuscript and the formulation of Wahl & Gerrish (2001) [2], who derived the fixation probability for beneficial mutations in periodically bottlenecked populations. We show that our per-cycle selection coefficient relates to the Wahl-Gerrish “selective advantage” (defined as the ratio of growth rates minus one) by a factor of *rτ*, where*r*is the wild-type growth rate and*τ*is the time between bottlenecks. We also demonstrate that the rate of establishment of beneficial mutations in the Wahl-Gerrish model is equivalent to the classical formula 2*sN*_*e*_*µ*, when parameters are appropriately interpreted.

#### S2.1 Notation and definitions

##### Wahl-Gerrish parameterization

In Wahl & Gerrish (2001), populations undergo exponential growth between periodic bottlenecks. The key parameters are:

- *N* _0_: population size immediately after a bottleneck
- *N* _*τ*_ : population size immediately before the next bottleneck (i.e., at time*τ*)
- *r*: wild-type Malthusian growth rate (units of time ^*−*1^)
- *τ*: time between bottlenecks
- *D*=*N* _0_*/N*_*τ*_ =*e* ^*−rτ*^ : dilution factor
- *s* _*w*_: selective advantage, defined as the ratio of mutant to wild-type growth rates minus one, such that the mutant growth rate is*r*(1 +*s* _*w*_)

With exponential growth, the population grows from *N* _0_ to*N* _*τ*_ =*N* _0_*e*^*rτ*^ during each cycle, followed by dilution back to*N* _0_.

##### Our parameterization

In our manuscript, we express quantities per growth cycle (dilution-to-dilution interval), treating the cycle as the natural unit of time:

- *N* _*b*_: bottleneck population size (≡*N* _0_)
- 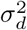 variance in descendant number per cycle
- 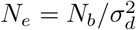 effective population size per cycle
- *s:* per-cycle selection coefficient (mean excess relative growth over one cycle)

##### S2.1.1 Relationship between selection coefficients

###### The Wahl-Gerrish selection coefficient

Consistent with the standard parameterization in the LTEE literature, Wahl & Gerrish define the selective advantage *s* _*w*_ via the ratio of Malthusian fitnesses, subtracted by one,

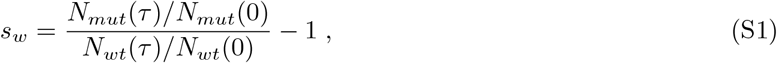

where*N* _*mut*_(*t*) and*N* _*wt*_(*t*) are the mutant and wild-type population sizes, respectively. In the Wahl-Gerrish model, the selective advantage solely depends on the growth rates,

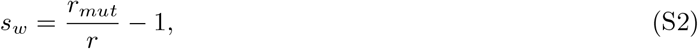

where*r* _*mut*_ and*r*are the Malthusian growth rates of mutant and wild-type, respectively.

###### Per-cycle selection coefficient

The per-cycle selection coefficient, which we denote*s*, measures the mean excess relative growth over one complete cycle, or the slope in the logit frequency of the mutant,

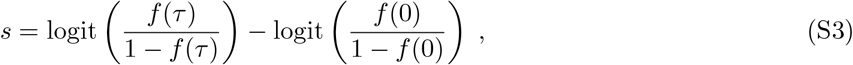

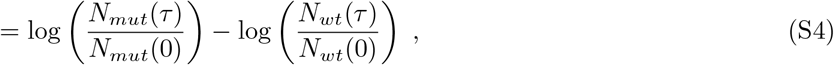

where*f*(*t*) is the mutant frequency.

For a mutant with growth rate*r* _*mut*_ =*r*(1 +*s* _*w*_), the per-cycle selective advantage is:

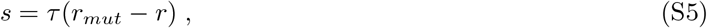

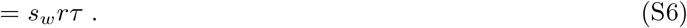

where*rτ*= ln(*N* _*τ*_ */N*_0_) = − ln*D*. For the LTEE with 1:100 daily dilutions,*rτ*= ln(100) ≈4.6, corresponding to about 6.64 generations per cycle.

#### S2.2 Effective population size in the Wahl-Gerrish Model

##### Time-dependent fixation probability

Wahl & Gerrish showed that a beneficial mutation (under weak selection, *s* _*w*_*rτ* ≪1) arising at time*t*during the growth phase (where *t* ∈ [0, *τ*]) has an approximate fixation probability of:

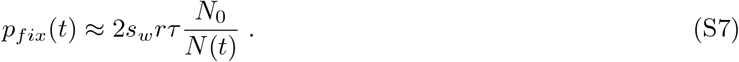

Using our definition of*s*,

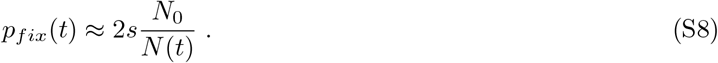

##### Effective population size

Wahl & Gerrish derived an effective population size for periodically bottle-necked populations (in units of number of generations; more precisely, in units of*e*-foldings):

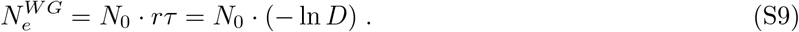

This effective population size captures the reduced efficacy of selection due to population bottlenecks. In our notation with per-cycle measurements, when the only source of descendant-number variance is the bottleneck itself (Poisson sampling with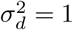), our effective population size (in units of number of cycles) is simply

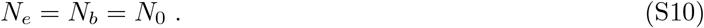

#### S2.3 Establishment rate

At time*t*during the growth phase, mutations arise at rate*µN*(*t*) per unit time, where *µ* is the per-cell mutation rate. The instantaneous establishment rate is:

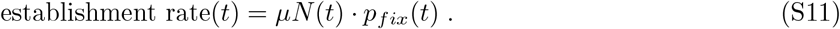

Substituting Eq. (S8):

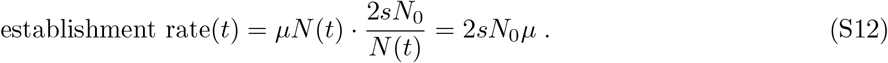

We see that the establishment rate is constant in time, independent of when during the cycle the mutation arises. This is because although mutations arising late in the cycle are more numerous (larger*N*(*t*)), they have proportionally lower fixation probability (smaller*N* _0_*/N*(*t*)), and these effects exactly cancel. This expression is equivalent to the establishment rate in classical population genetics models [3].

### S3 Theory: Noisy Growth Traits

#### S3.1 Variation in lag times

We consider a population of asexually reproducing microbes that is transferred to a new environment. After an initial lag time, individuals start growing exponentially at a growth rate*r*. The culture is allow to grow for a total duration*T*.

##### S3.1.1 Exponential distribution of lag times

Suppose that the lag time for each individual,*t* _*lag*_, is an exponential random variable,

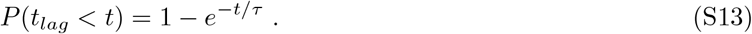

The total population size of a clone that starts as a single individual at time 0 will be*N*= exp[*r*(*T*− *t* _*lag*_)] at time*T*. It follows that,

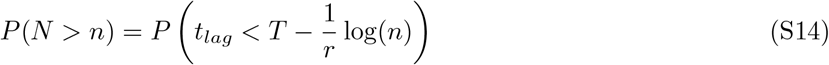

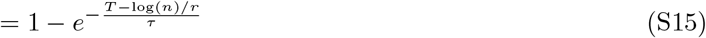

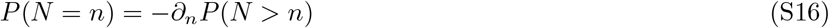

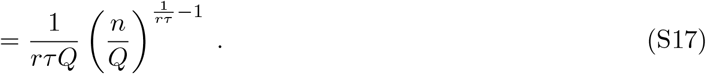

We define *Q*=*e* ^*rT*^ as the maximum size of a clone (i.e. if it starts growing immediately;*t* _*lag*_ = 0). The expression is valid for *n < Q* and assuming that *P*(*t* _*lag*_ *> T*) ≈ 0. We see that the offspring number distribution*P*(*N*=*n*) is a power-law, with an exponent that is always larger than 1. For notational simplicity, we define a compound parameter*γ*= − 1*/*(*rτ*). We can now calculate the variance of clone sizes,

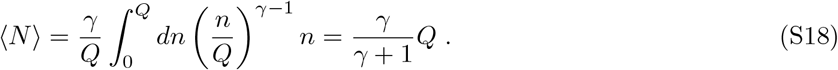

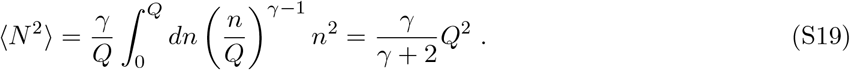

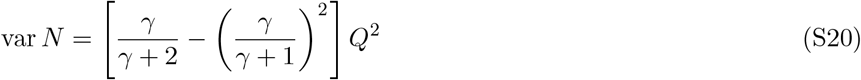

In a serial dilution regime, the population size at time*T*will be diluted down by a factor of ⟨*N*⟩, so that each clone will start again with one individual, on average. Thus, the effective additional descendant number variance across cycles is,

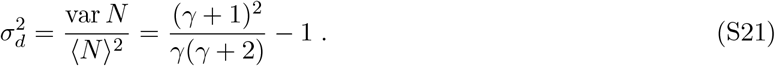

When 1*/γ*≫1, we see that the descendant number variance increases linearly with the lag time mean/standard deviation,*σ* ^2^ ∝*τ*.

##### S3.1.2 Gamma distribution of lag times

Following closely the derivation of the prior section, we now consider the case that the lag time for each individual,*t* _*lag*_, is a gamma random variable,

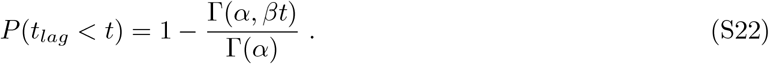

Here Γ(*·*) is the gamma function, and Γ(*·, ·*) is the upper incomplete gamma function. In this parameterization, the mean of the gamma distribution is*µ*=*α/β*and the variance is*v*=*α/β* ^2^. As before, the total population size of a clone that starts as a single individual at time 0 will be*N*= exp[*r*(*T* −*t* _*lag*_)] at time*T*. It follows that,

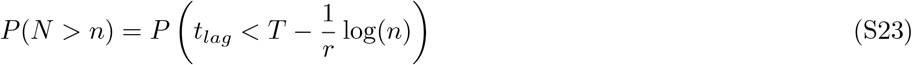

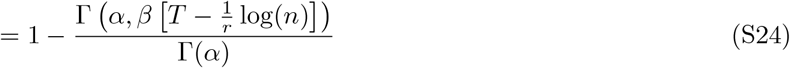

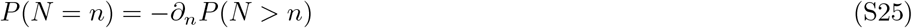

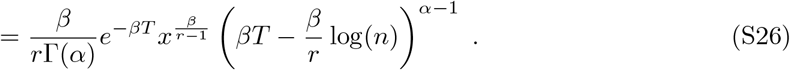

Again, we define*Q*=*e* ^*rT*^ as the maximum size of a clone (i.e. if it starts growing immediately;*t* _*lag*_ = 0). The expression is valid for*n < Q*and assuming that*P*(*t* _*lag*_ *> T*)≈0. We calculate the moments as before,

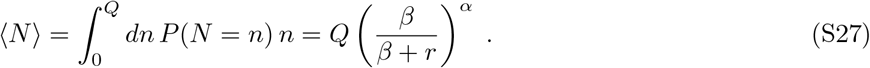

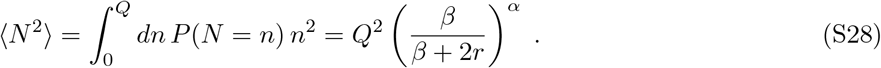

In a serial dilution regime, the population size at time*T*will be diluted down by a factor of ⟨*N*⟩, so that each clone will start again with one individual, on average. Thus, the effective additional descendant number variance across cycles is,

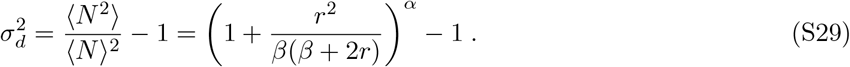

#### S3.2 Variation in growth rates

We consider the case that the exponential growth rate is noisy. Again, we consider an asexual population of microbes that are growing in exponential phase for some time*T*. Using continuous time models, we take the time to split into two daughter cells as a random variable; we neglect mother-daughter correlations.

##### S3.2.1 Exponential distribution of doubling times

The simplest case of noisy growth rates is when doubling times are exponentially distributed with a mean parameter*τ*, as division is memoryless (i.e. a Poisson process). This problem can be formulated as a Galton-Watson process [4], which gives first and second moments of the population sizes,

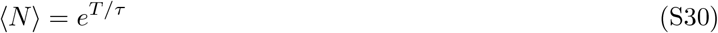

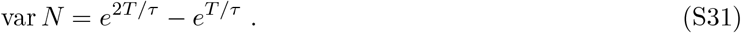

As before, in a serial dilution regime, the population size at time*T*will be diluted down by a factor of ⟨*N* ⟩, so that each clone will start again with one individual, on average. Thus, the effective additional descendant number variance across cycles is,

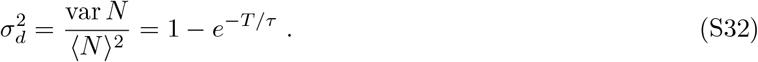

##### S3.2.2 Gamma distribution of doubling times

We now assume that doubling times are distributed according to a gamma distribution. We treat the process as a classical Bellman-Harris process, and obtain asymptotic solutions for the moments [5]. As the total time becomes large,*T*→ ∞, the mean number of individuals can be approximated by,

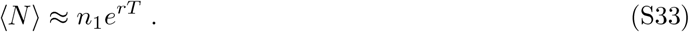

Here,*r*is the Malthusian parameter, which is the solution to the integral,

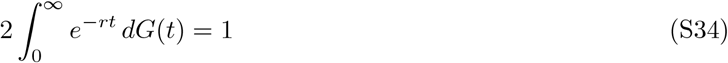

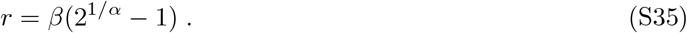

The function*G*(*t*) is the cumulative distribution function of the gamma distribution,

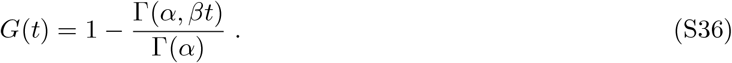

The prefactor*n* _1_ is given by the following integral,

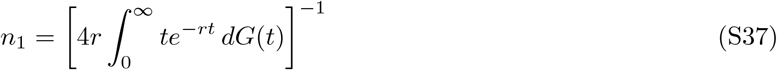

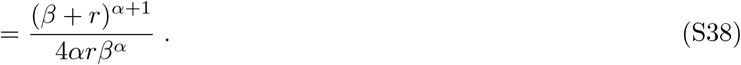

Similarly, the second moment can be asymptotically approximated by,

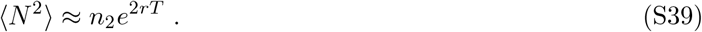

The prefactor*n* _2_ is given by,

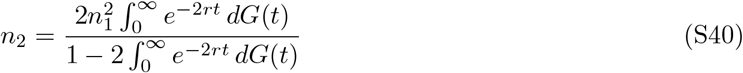

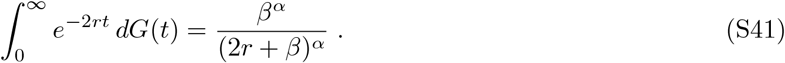

Finally, the effective additional descendant number variance is given by,

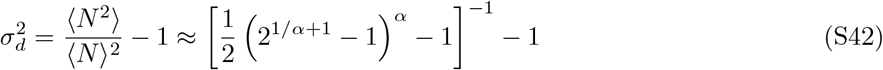

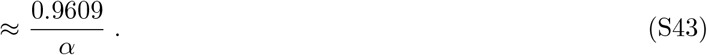

We see that the effective descendant number variance does not depend on*T*or*β*. It only depends on*α*, which is related to the coefficient of variation of the gamma distribution,*α*=*µ* ^2^*/σ*^2^. The effective descendant number variance is well approximated by a simple inverse relationship to*α*.

---

**Figure S1.**
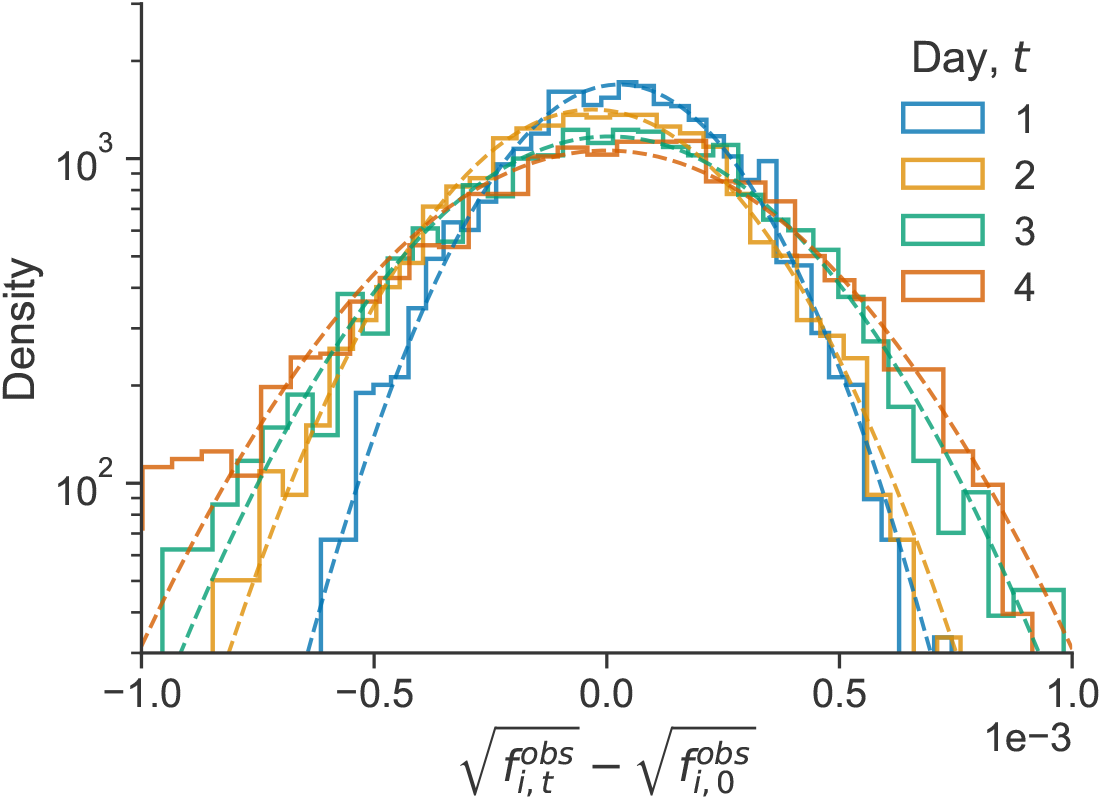
Variance in allele frequency trajectories increases over time(related to Figures 1D and 1E). Transition densities of barcode frequencies for the REL606 library, replicate 1. Here, we compute the transition densities in reference to the frequency at time point 0. We see that the variance increases over time, which is expected given that variance due to genetic drift accumulates over time. Dotted lines represent Gaussian fits to the empirical densities.

**Figure S2.**
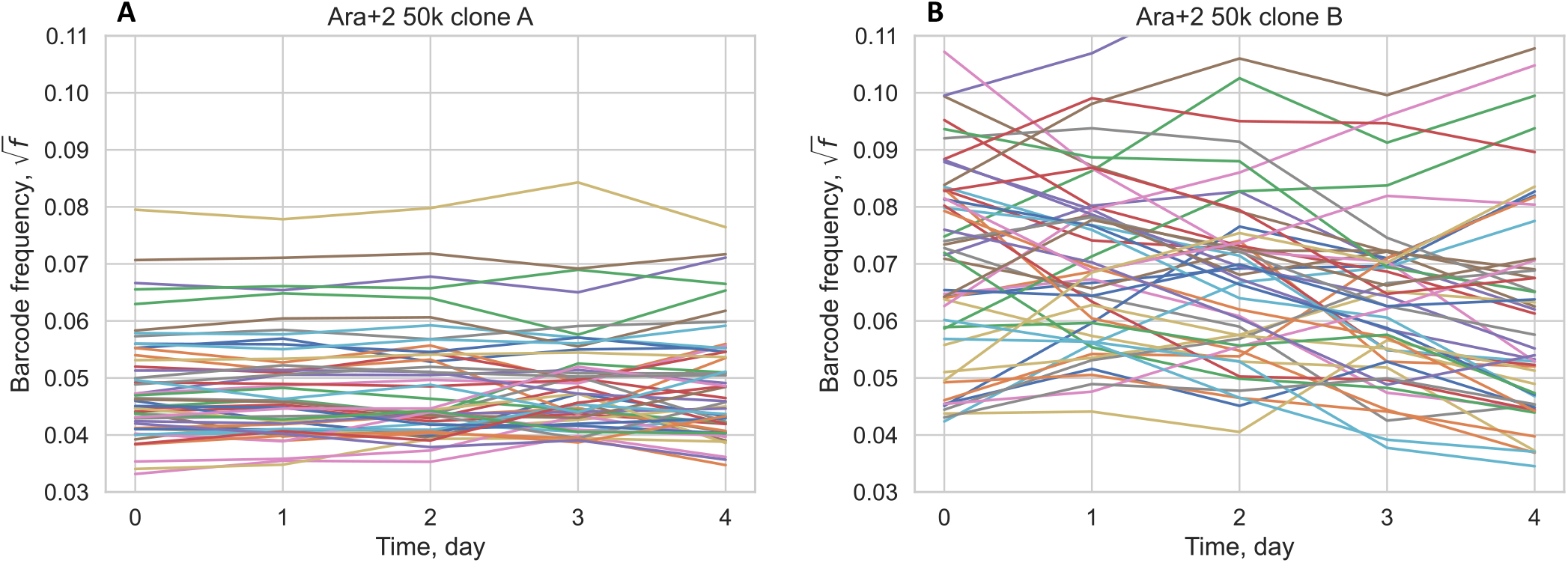
Ara+2 50k barcode frequency trajectories.We show example square-root transformed barcode frequency trajectories for clones. (**A**)*A* and (**B**) *B*. We randomly sampled 50 barcodes from each clone, both from biological replicate 1. Clone*B*visibly has stronger fluctuations compared to clone *A*.

**Figure S3.**
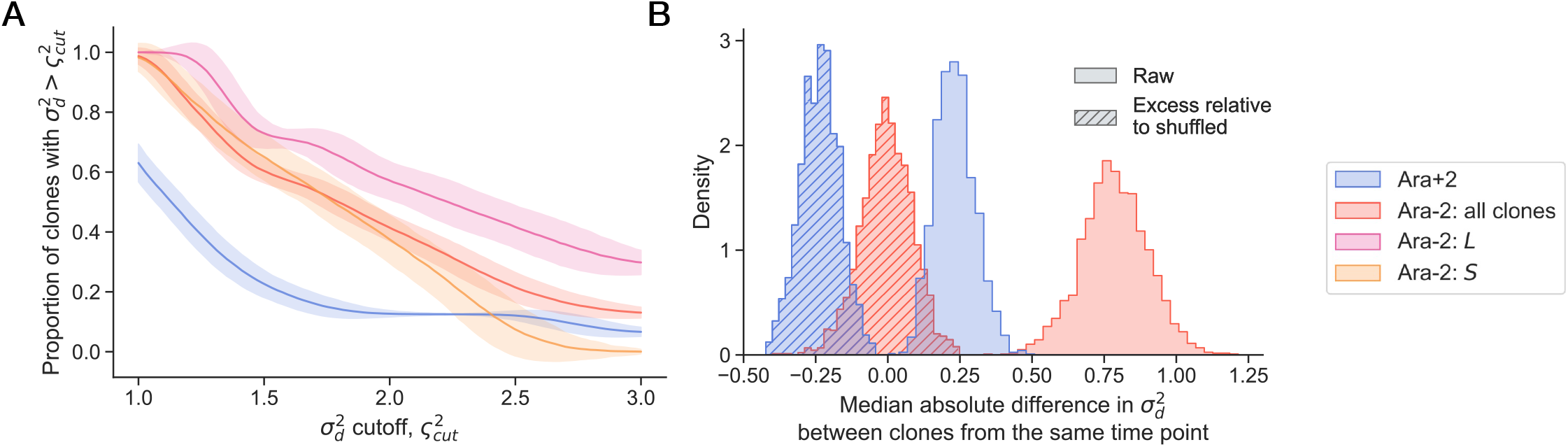
Additional patterns of the strength of genetic drift across lineages. (**A**) We computed the proportion of clones within a group that have an inferred descendant number variance larger than a cutoff value. Error bars represent the posterior standard deviation. (**B**) We sought to understand if clones from the same evolutionary time point displayed similar strengths of genetic drift. For each pair of clones within a lineage (Ara+2 or Ara-2), we compute the absolute difference between their descendant number variances, and then compute the median across all clone pairs within the line (solid posterior densities). We see that the median difference in 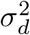 is generally larger in the Ara-2 line, compared with the Ara+2 line. This could reflect that fact that Ara+2 clones at the same evolutionary time point are more genetically related to each other than Ara-2 clones at the same time point. We were curious to see if clones at the same evolutionary time point were more or less similar to each other than the random expectation. Thus, we shuffled the identity of the clones, repeated the process, and subtracted the shuffled differences from the raw differences (circle hatched posterior densities). We see that the excess difference in 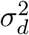 relative to the shuffled data is centered around zero for Ara-2, but it is below zero for Ara+2. This indicates that Ara+2 clones at the same evolutionary time point are generally more similar to each other than would be expected if there was no time point stratification. On the other hand, we do not see the same similarity between Ara-2 clones at the same time point.

**Figure S4.**
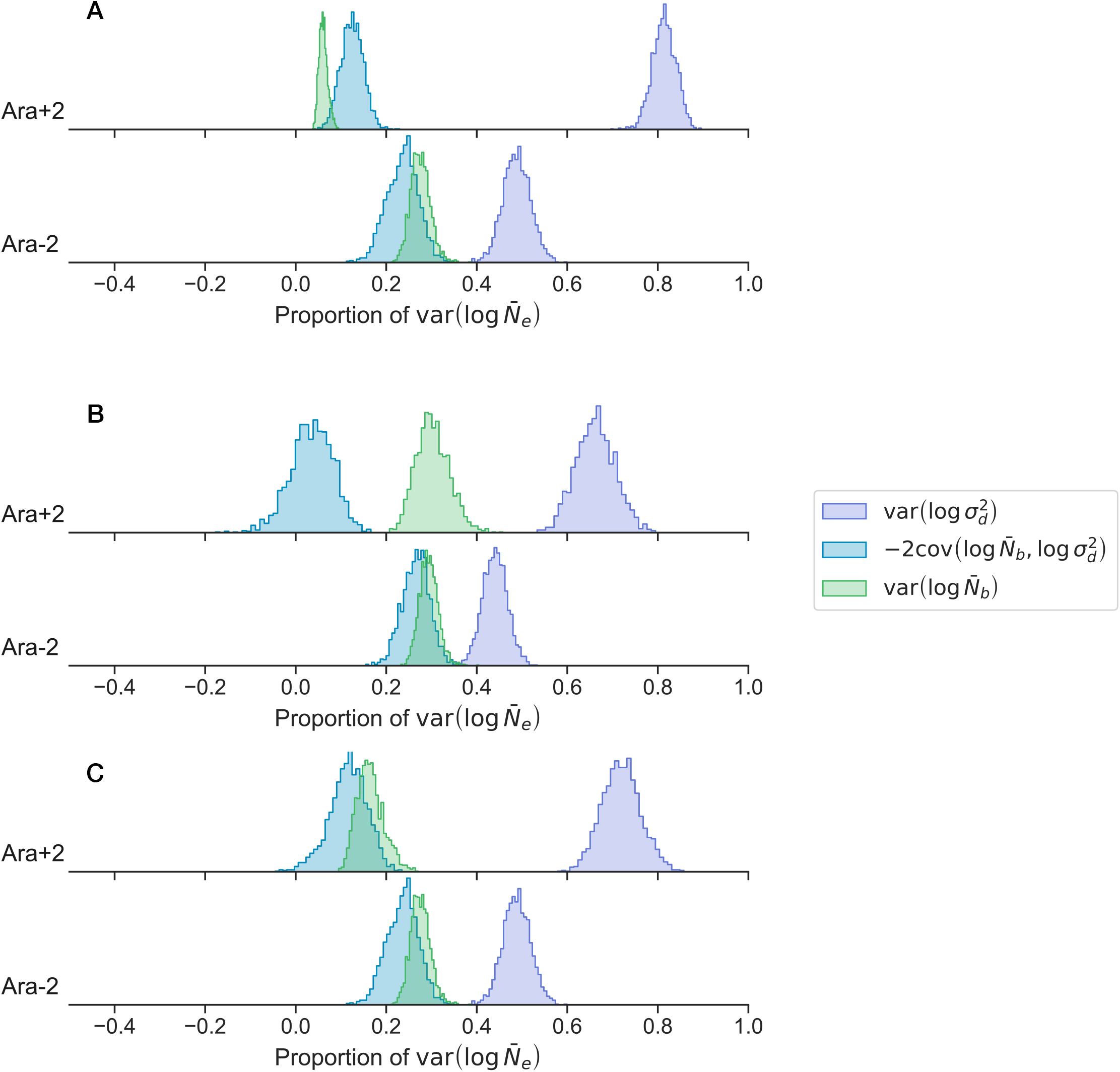
Contributions to var log 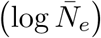. Same as Figure 4E, except (**A**) REL606 is excluded from the calculation, (**B**) the Ara+2 50k B clone is excluded (i.e. the outlier clone with the large 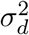 value), or (**C**) both REL606 and the Ara+2 50k B clone is excluded.

**Figure S5.**
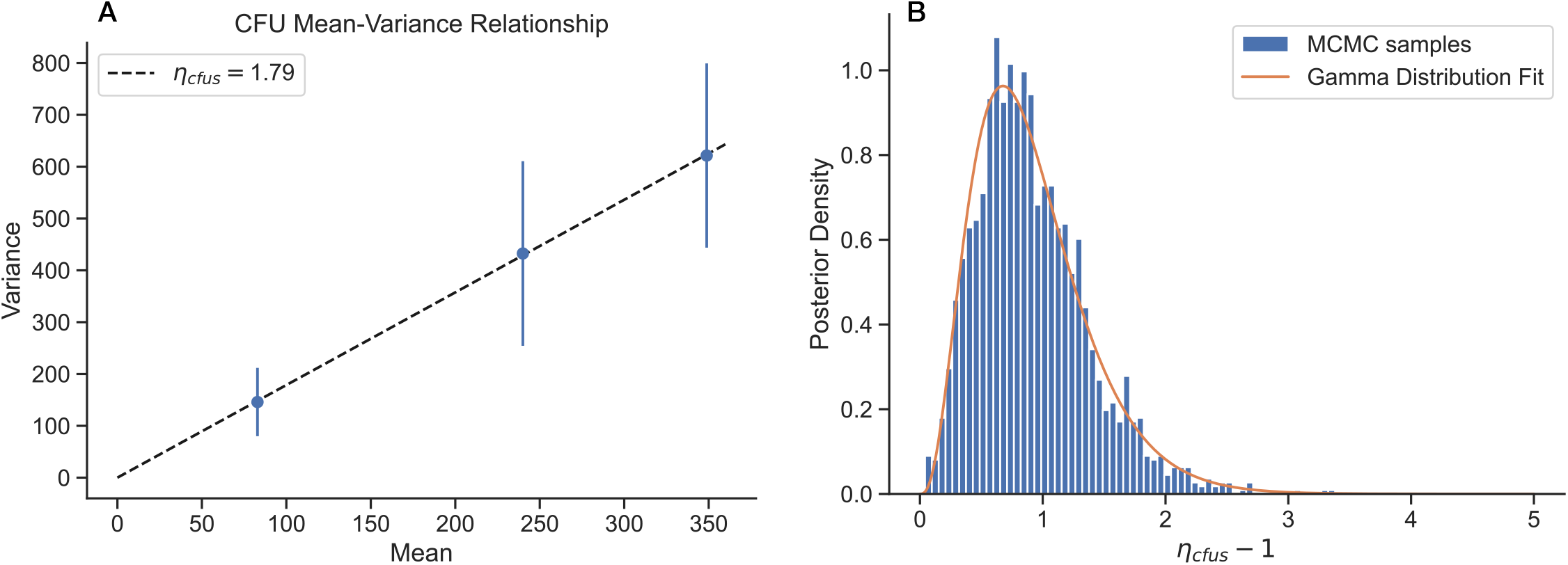
Measurement of CFU errors. (**A**) Using three different cultures of REL606 at different cell densities, we diluted and plated the cultures on agar plates as 16 independent technical replicates. We then computed the mean and variance across technical replicates for each culture. We see a linear mean-variance relationship. (**B**) Using a Bayesian inference framework, we obtained an estimate for the marginal posterior of the overdispersion parameter*η* _*cfus*_ (see Methods subsection “Bayesian inference of genetic drift parameters”).

**Figure S6.**
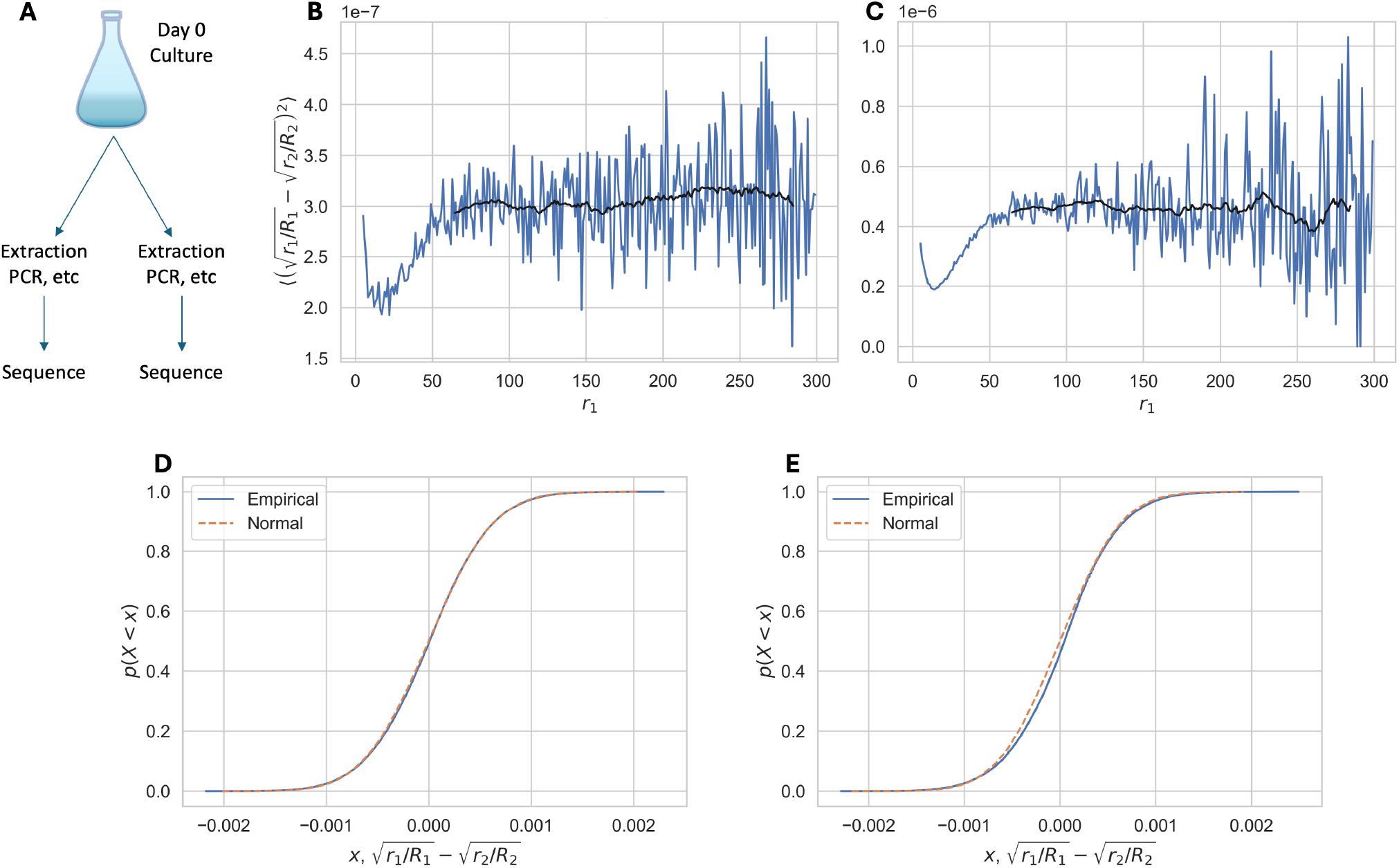
BarSeq technical replicates reveal patterns of measurement noise. (**A**) A single single culture of barcoded clones was split into two subcultures, and all sample processing steps (genomic DNA extraction, barcode amplification, sequencing) were done independently on the two cultures. (**B-C**) We quantified the average variance of square-root barcode frequencies, as a function of the read count of technical replicate 1,*r* _1_, for the (**B**) 6.5k*L*and (**C**) 6.5k*S*barcoded libraries. Blue lines represent raw data, black lines represent a rolling average. We see that the variance of the square-root transformed data is approximately stable for reads counts greater than about 50. (**D-E**) We found that the distribution of difference in frequencies between technical replicates was well approximated by a Gaussian distribution for both the (**D**) 6.5k *L* and (**E**) 6.5k *S* barcoded libraries.

**Figure S7.**
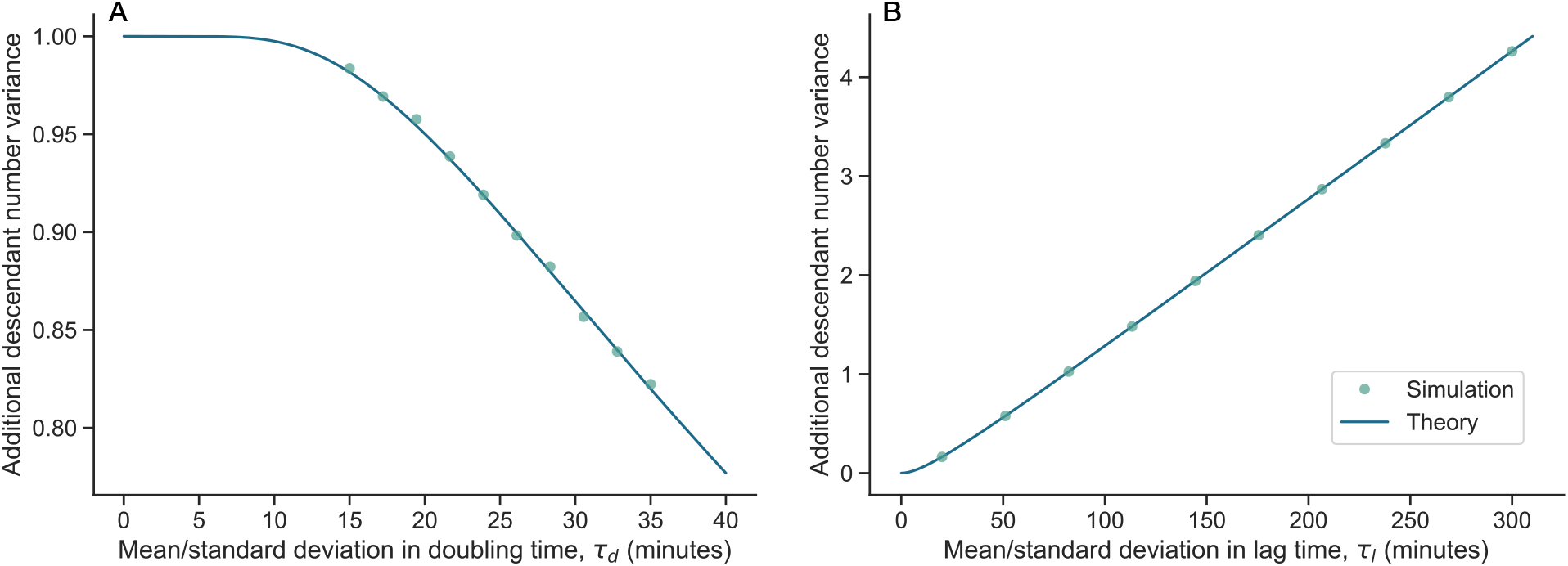
Physiological causes of changes to descendant number variance.Related to Figure 5A-B. We treat both (**A**) doubling times and (**B**) lag times as exponentially distributed random variables. Solid lines represent analytical results presented in sectionS3, and circles represent simulation results.

**Figure S8.**
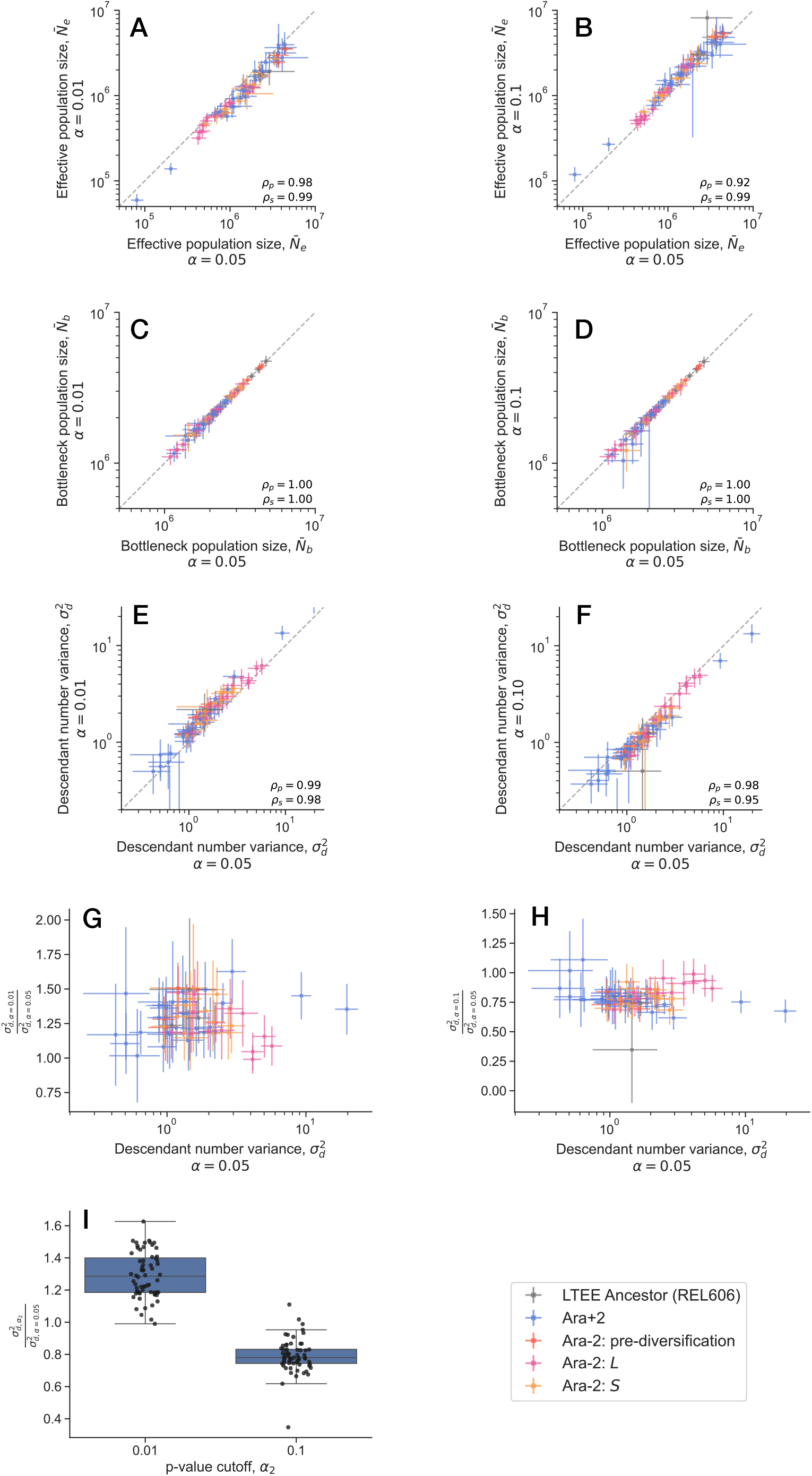
Results are robust to outlier detection threshold(*continued on next page*). We ran our pipeline to infer genetic drift parameters on our data, while varying the outlier detection cutoff (α), comparing the resulting inferred parameters to those presented in the main text, whereα= 0.05 (Figure 3). We usedα= 0.01 andα= 0.1. (A-F) We find that the inferred 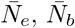 and 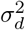 are all robust to changes inα. (G-I) It appears that lower αgenerally results in a higher inferred 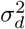, while higherαgenerally results in a lower inferred 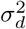. However, the magnitude of this effect is relatively small, typically changing the inferred value by roughly±25%.

**Figure S9.**
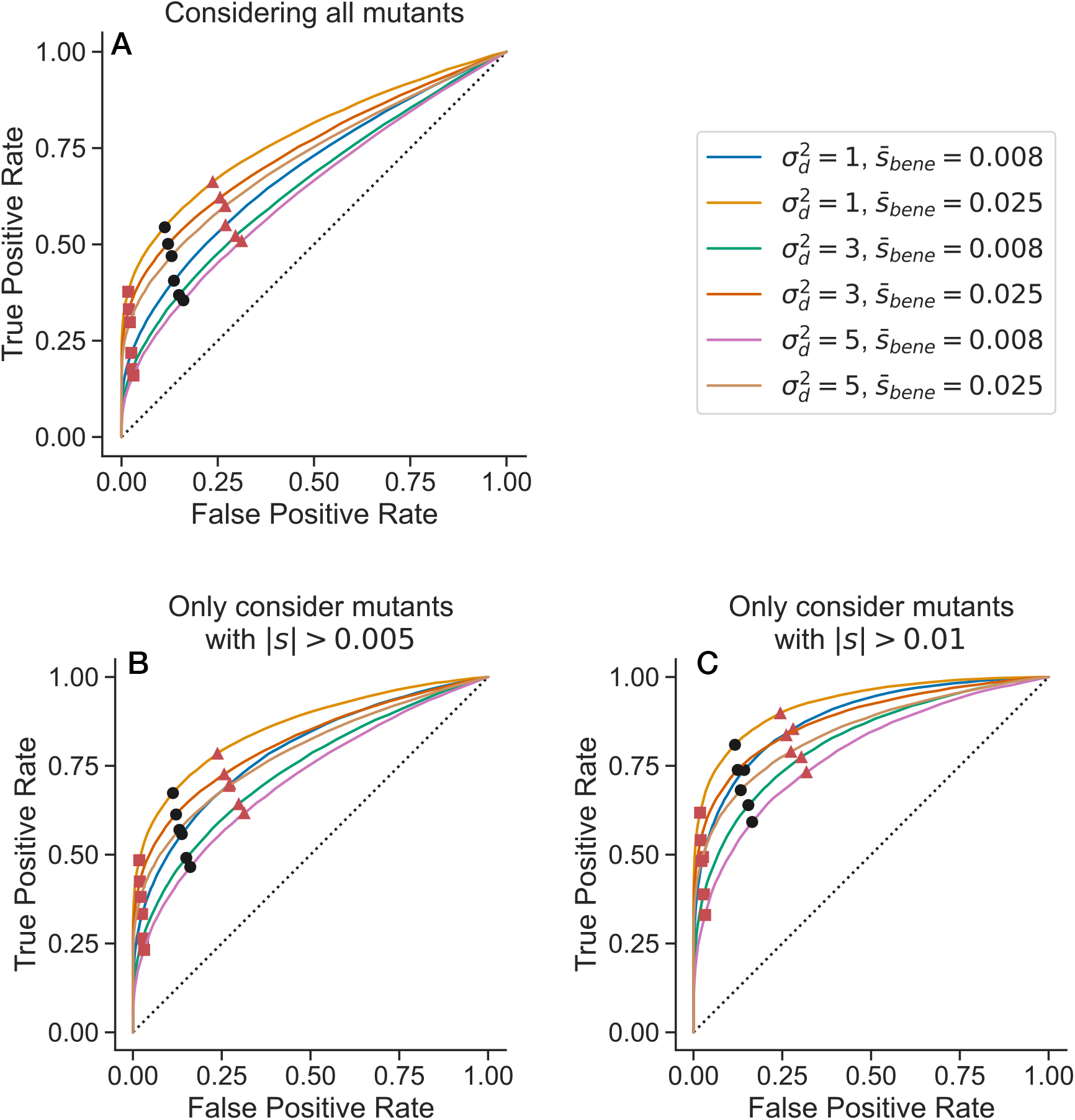
Receiver operating characteristic (ROC) curves for outlier detection. (**A**) Across all of our simulated data sets with contaminated non-neutral barcodes (see Methods subsection “Simulations of the data generating process and inference pipeline”), as we vary the threshold*p*value (*α*) to call a lineage as non-neutral, we compute the true and false positive rate that the lineage is actually non-neutral. (**B-C**) We perform the same computation, but testing our ability to distinguish larger fitness effect mutants from neutrality, considering (**B**) | *s* |*>*0.005 or (**C**) |*s*| *>*0.01. Black dots represent the point where*α*= 0.05; red triangles represent where*α*= 0.1; red squares represent where*α*= 0.01.

**Figure S10.**
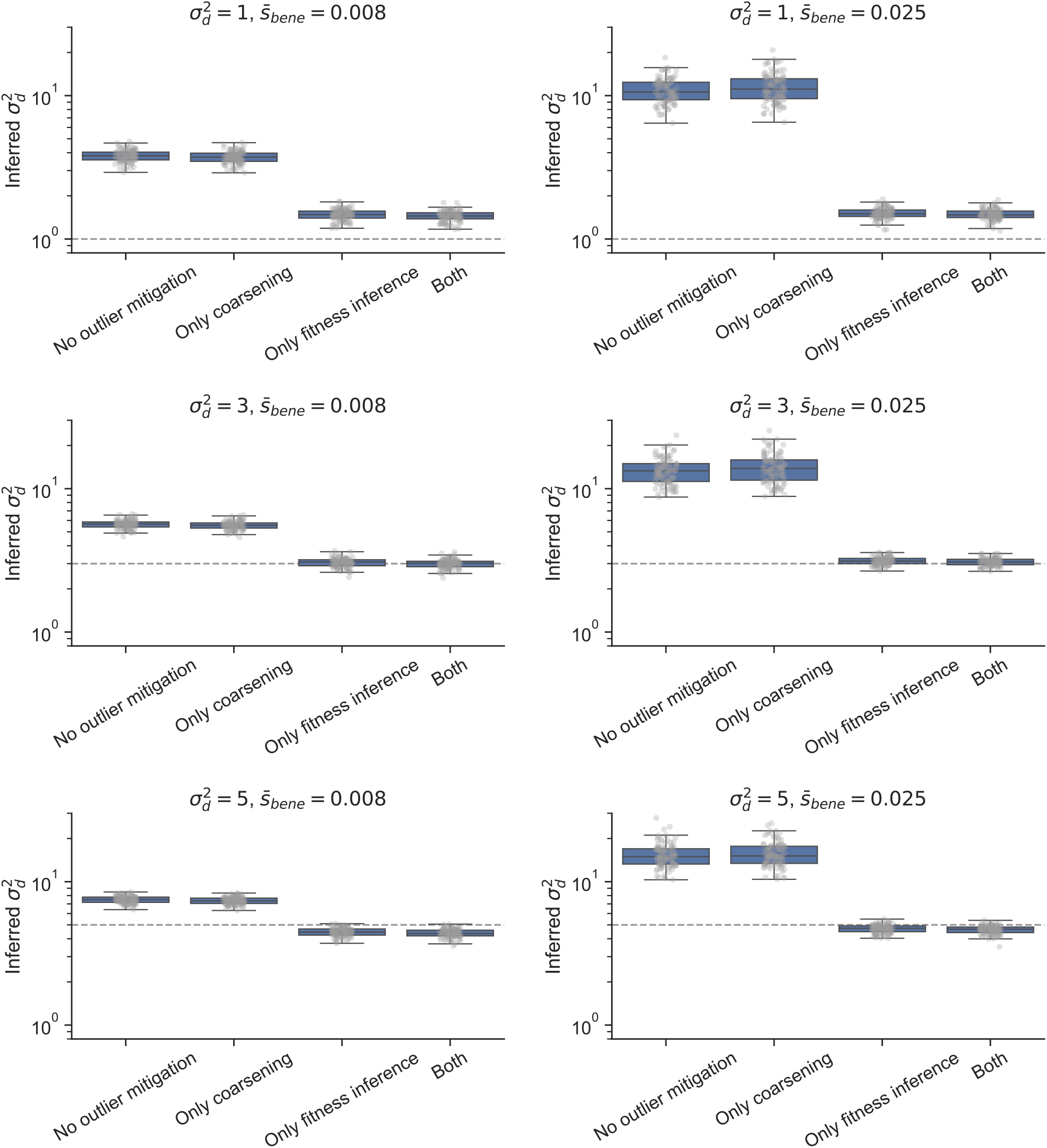
Simulation results for 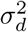 inference. We ran simulations of the data generating process and drift inference pipeline to validate if our method is able to recover the true values of drift parameters, even in the case of contaminating non-neutral barcodes (see Methods subsection “Simulations of the data generating process and inference pipeline”). We varied the true 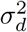 and the mean of the beneficial side of the distribution of fitness effects,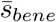. We compare two outlier mitigation measures, alone and together. We find that fitness inference and exclusion of putatively non-neutral barcodes (here at*α*= 0.05) is particularly crucial to recover nearly unbiased estimates of 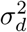 across conditions. Likelihood coarsening only regularizes the estimates slightly.

**Figure S11.**
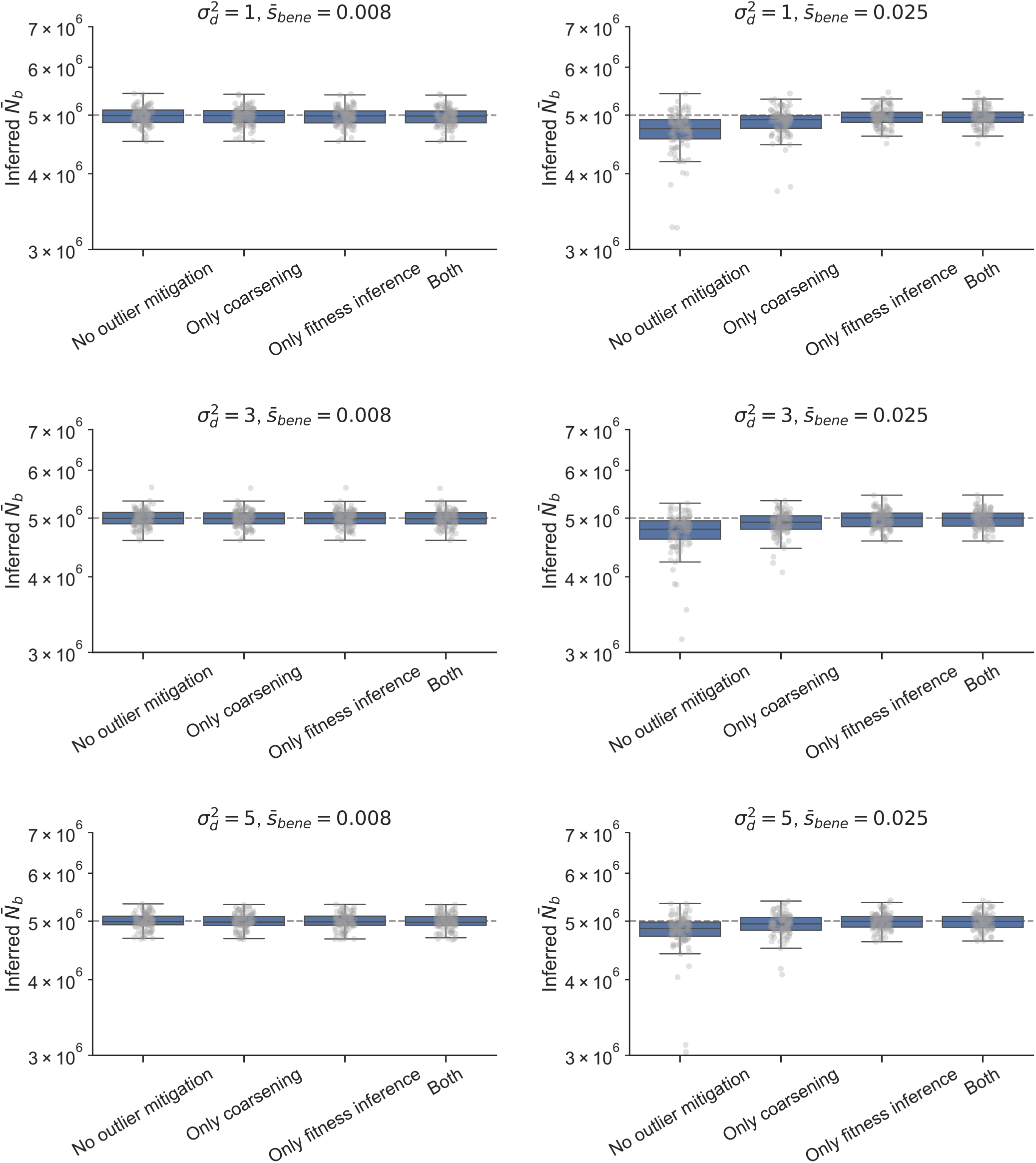
Simulation results for 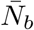 inference. See caption of Figure S10–here we show the results for 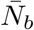 inference instead of 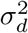 inference. Generally 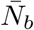 inference is robust to the presence of outlier barcodes.

**Figure S12.**
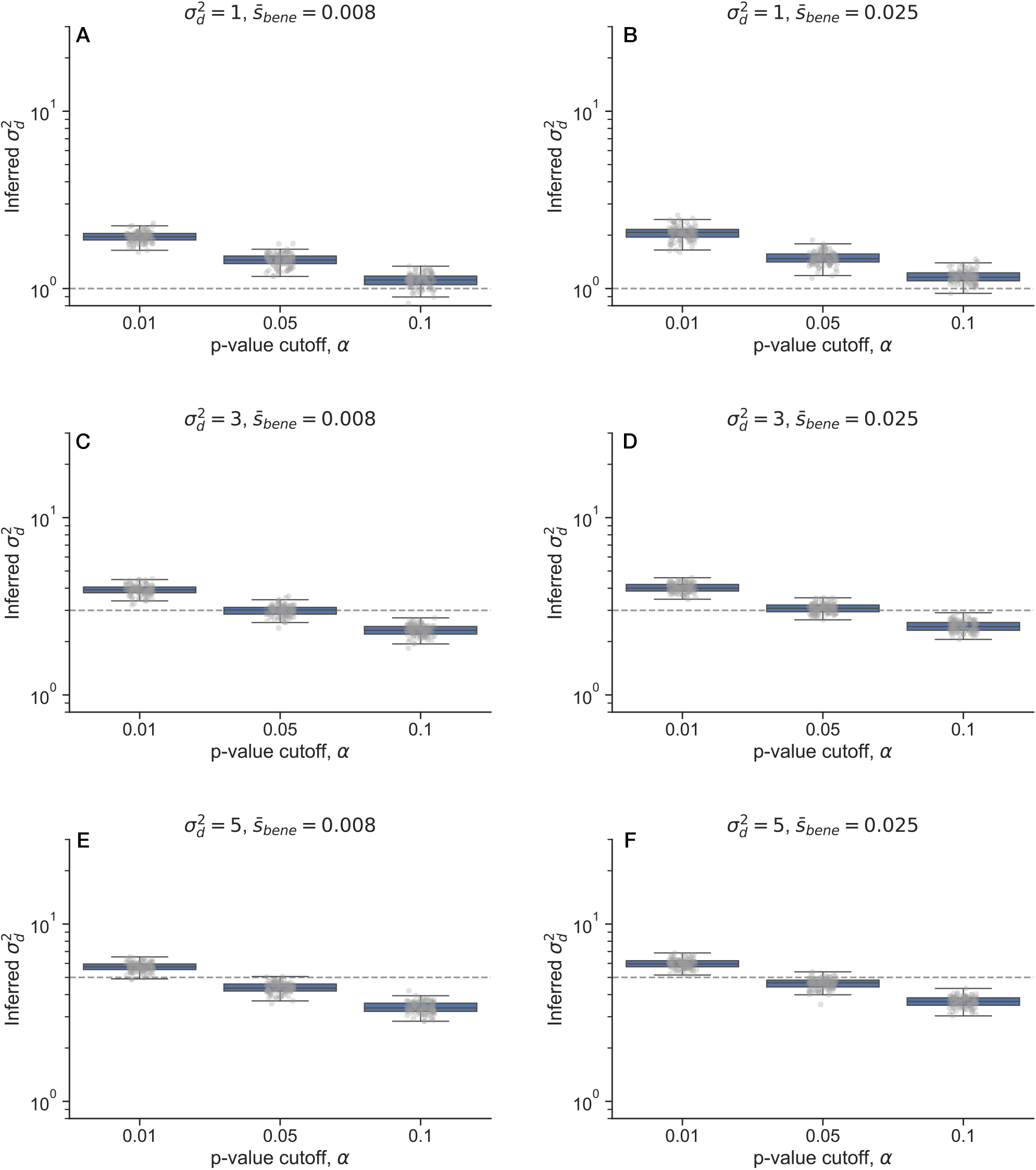
The effect of varying the outlier detection cutoff on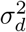 in simulations. We ran simulations of the data generating process and drift inference pipeline to validate if our method is able to recover the true values of drift parameters, even in the case of contaminating non-neutral barcodes (see Methods subsection “Simulations of the data generating process and inference pipeline”). We varied the true 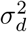 and the mean of the beneficial side of the distribution of fitness effects,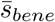 . We varied the p-value cutoff (*α*) for the likehood-based outlier detection. We see that *α*= 0.01 and *α*= 0.1 systematically over- and under-estimate 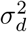 more often than the *α*= 0.05 cutoff, putatively by being too conservative or liberal, respectively, in excluding barcodes.

**Figure S13.**
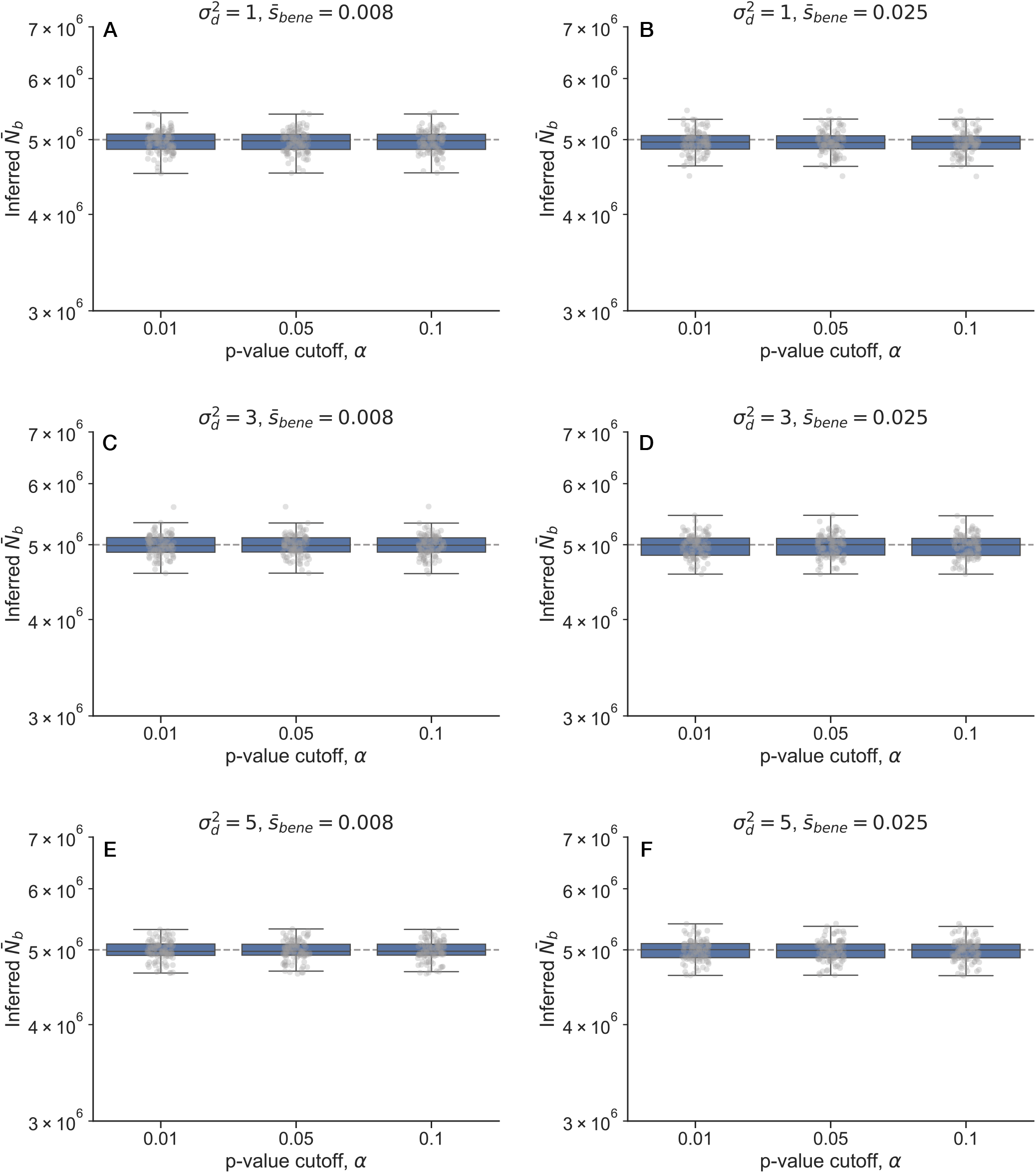
The effect of varying the outlier detection cutoff on 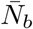 in simulations. See caption of Figure S12–here we show the results for 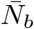 inference instead of 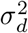 inference. Generally 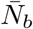 inference is robust to the presence of outlier barcodes.

**Figure S14.**
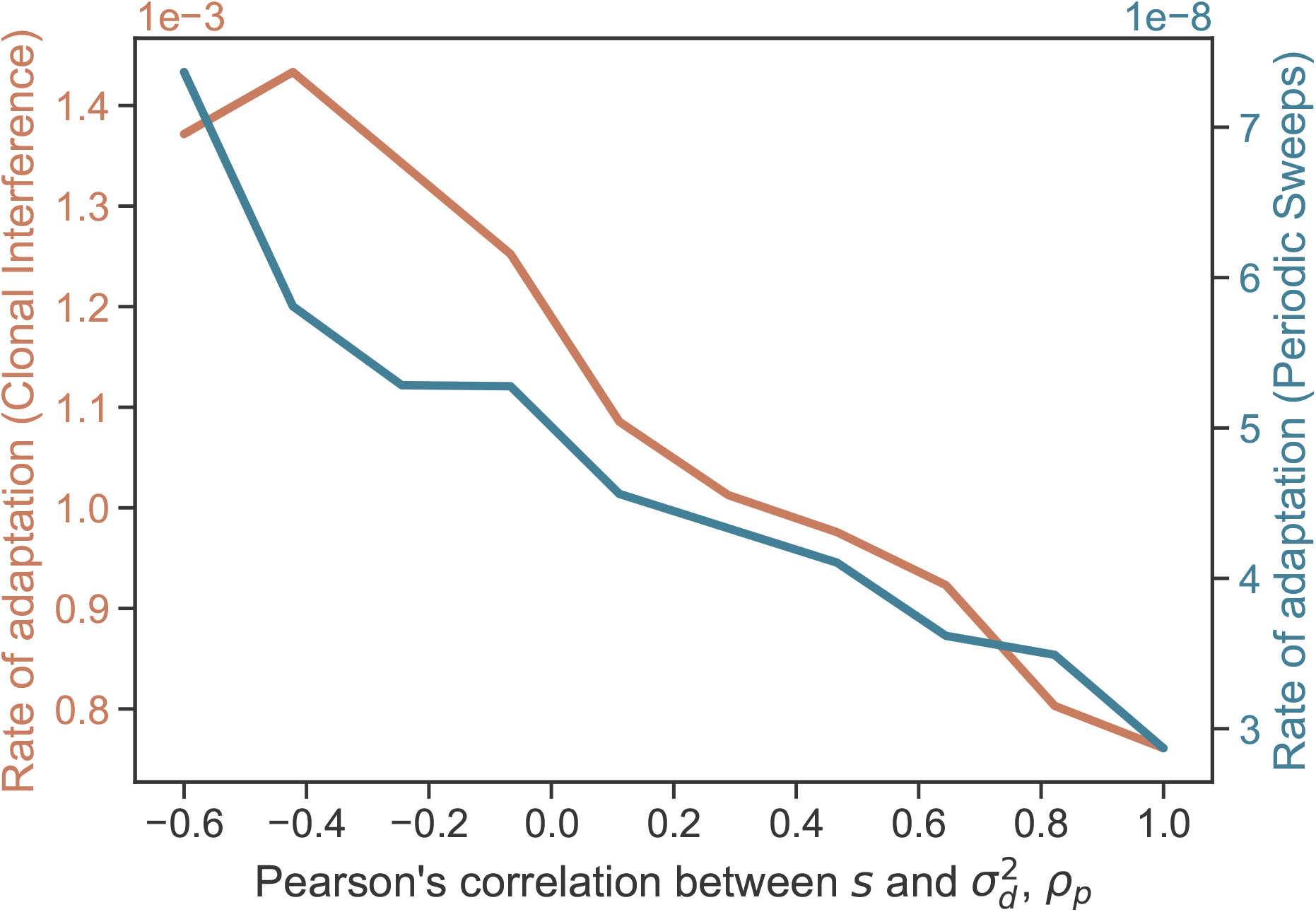
Effect of correlated mutation distributions on rate of adaptation. Related to Figure 5C-F in the main text. In both regimes, a stronger positive correlation between *s* _*mut*_ and 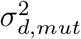is associated with a lower rate of adaptation (i.e. the average rate of fitness change over time). Here,*ρ* _*p*_ represents Pearson’s correlation coefficient of*p*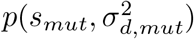.

**Figure S15.**
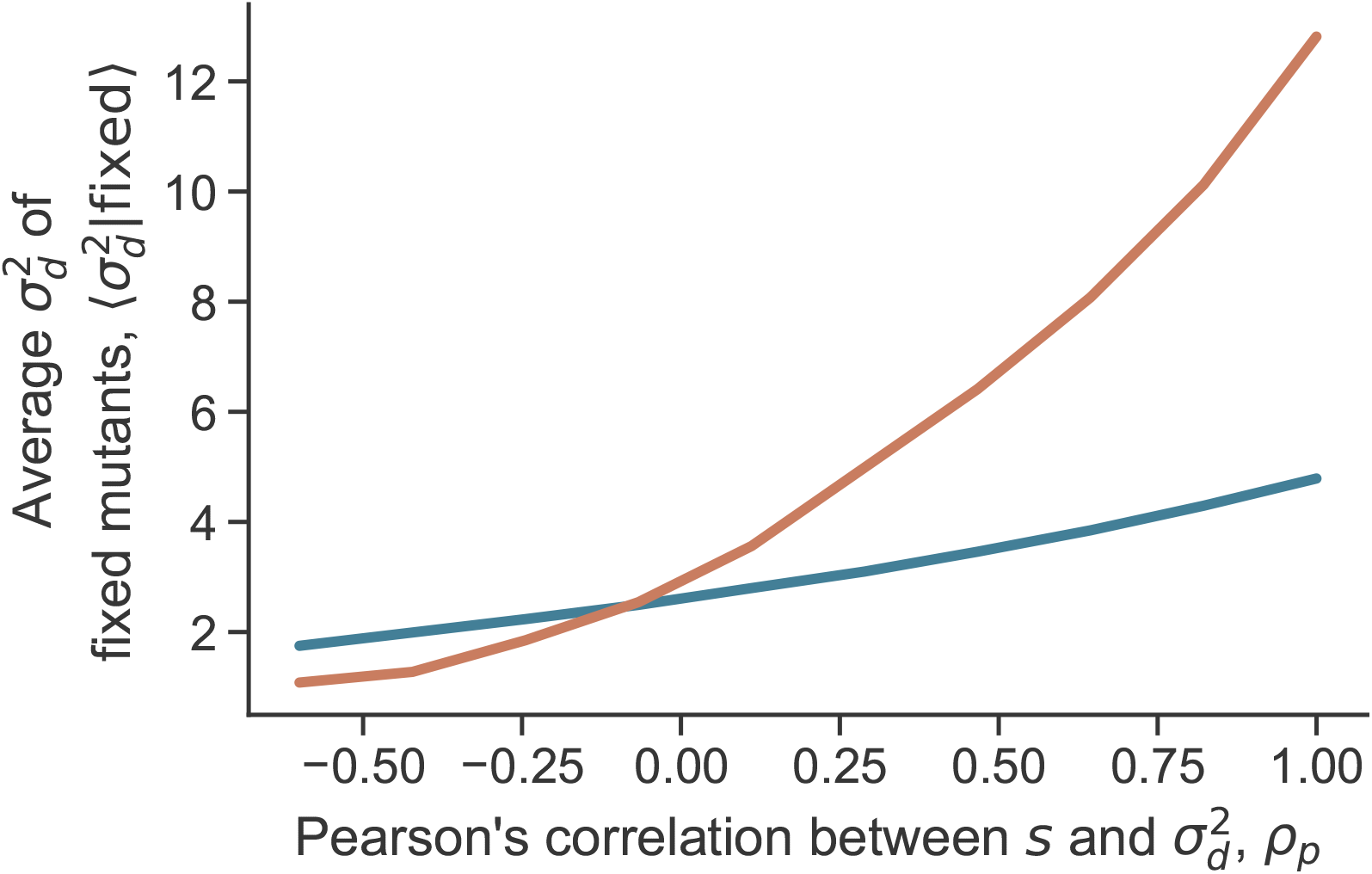
Effect of correlated mutation distributions on drift evolution. Related to Figure 5E in the main text. In both regimes, a stronger positive correlation between*s* _*mut*_ and 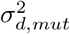 is associated with a higher 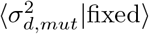. Here,*ρ* _*p*_ represents Pearson’s correlation coefficient of*p*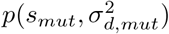.

**Figure S16.**
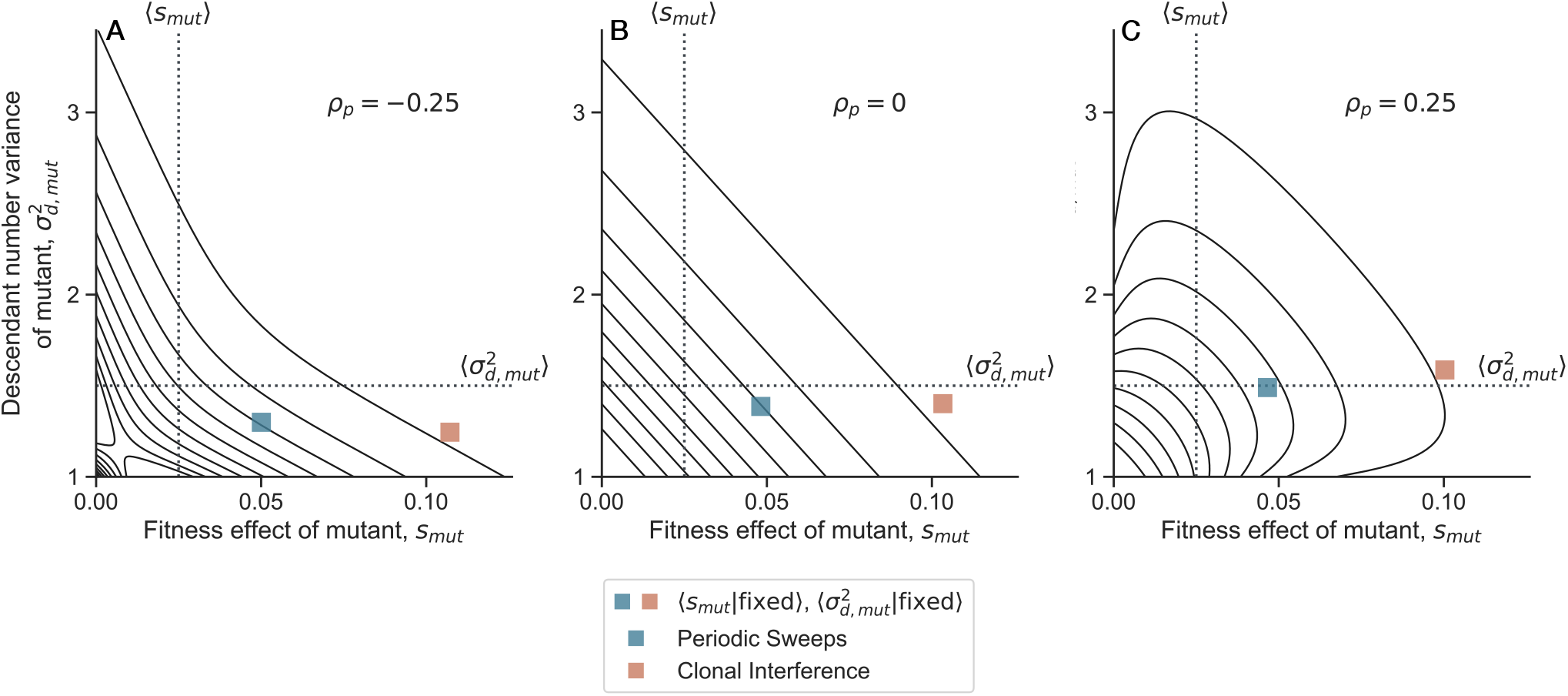
Joint distribution of fitness and drift effect–Gumbel’s bivariate exponential distribution. Related to Figure 5C-D in the main text, where here we use Gumbel’s bivariate exponential distribution instead of the Gaussian copula construction.

**Figure S17.**
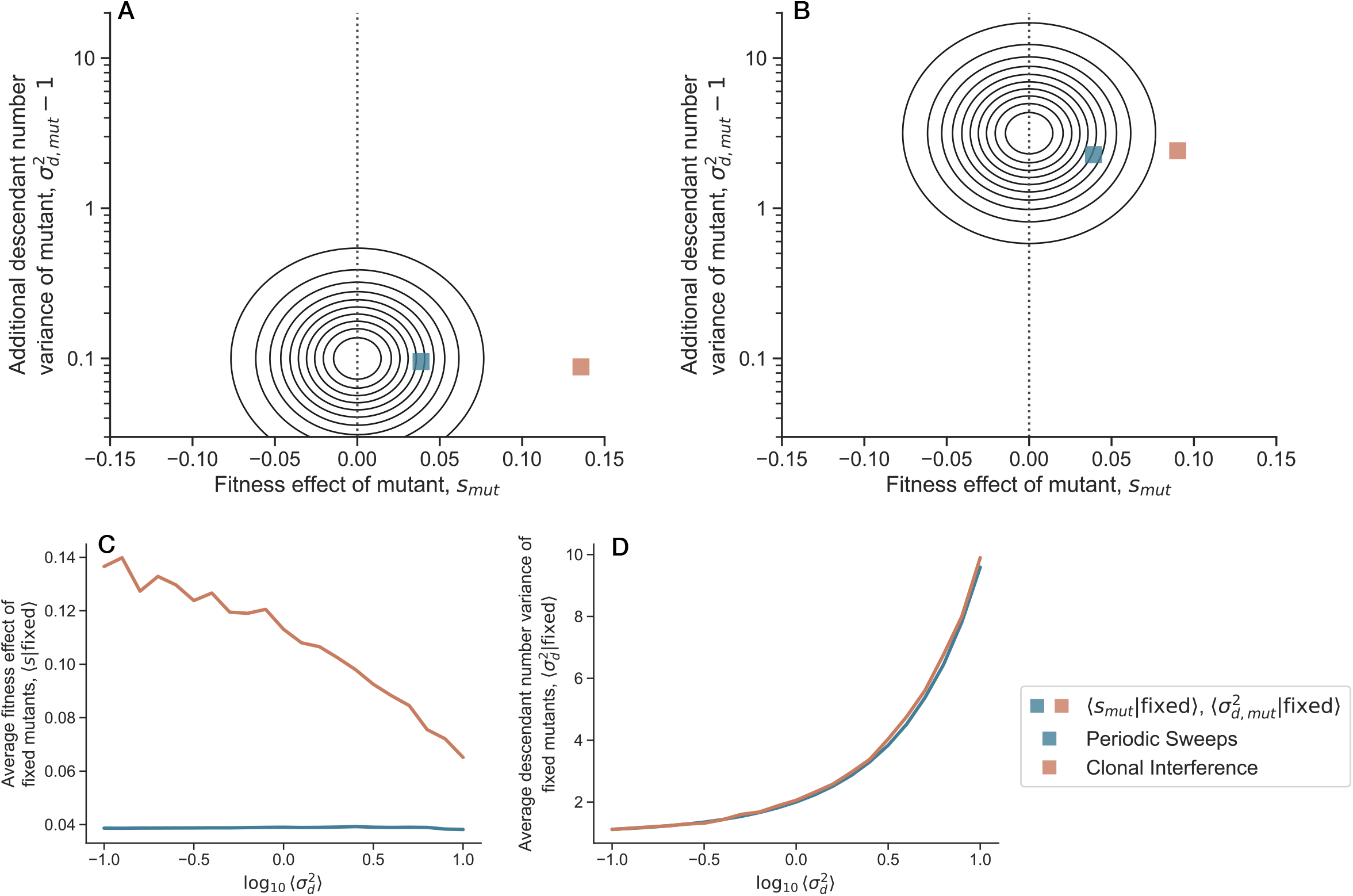
Joint distribution of fitness and drift effect–uncorrelated distribution. Related to Figure 5C-F in the main text, where here *s* _*mut*_ and 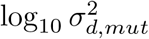 are drawn from an uncorrelated Gaussian distribution. As we increase the mean of 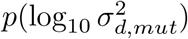, we see that the average fitness effect of fixed mutations stays constant in the periodic sweeps regime, while decreasing in the clonal interference regime.

## Notes

### Competing Interest Statement

The authors have declared no competing interest.

### Summary of Updates

1.We revised main text figures: a.Figure 1: Added panel A; revision of panel F. b.Figure 2: Added panels A and B. c.Figure 4: Added panel C; moved former panels C and D to SI d.Figure 5: Added panels E-F, revision of panels C-D. 2.We have shown that our results our robust to changes in the outlier detection threshold (Figure S8), and we performed additional simulations to probe the effect outlier detection threshold and further validate our choice of threshold (Figure S12-S13).

